# Roles of proteins containing immunoglobulin-like domains in the conjugation of bacterial plasmids

**DOI:** 10.1101/2021.07.22.453378

**Authors:** Mário Hüttener, Jon Hergueta, Manuel Bernabeu, Alejandro Prieto, Sonia Aznar, Susana Merino, Joan Tomás, Antonio Juárez

## Abstract

Horizontal transfer of bacterial plasmids generates genetic variability and contributes to the dissemination of the genes that enable bacterial cells to develop antimicrobial resistance (AMR). Several aspects of the conjugative process have long been known, namely, those related to the proteins that participate in the establishment of cell-to-cell contact and to the enzymatic processes associated with the processing of plasmid DNA and its transfer to the recipient cell. In this work, we describe the role of newly identified proteins that influence the conjugation of several plasmids. Genes encoding high-molecular-weight proteins that contain one or several immunoglobulin-like domains (Big) are located in the transfer regions of several plasmids that usually harbor AMR determinants. These Big proteins are exported to the external medium and target two extracellular organelles: the flagella and the conjugative pili. The plasmid-encoded Big proteins facilitate conjugation by reducing cell motility and facilitating cell-to-cell contact by binding both to the flagella and to the conjugative pilus. They use the same export machinery as that used by the conjugative pilus components. In the examples characterized in this paper, these proteins influence conjugation at environmental temperatures (i.e., 25°C). This suggests that they may play relevant roles in the dissemination of plasmids in natural environments. Taking into account that they interact with outer surface organelles, they could be targeted to control the dissemination of different bacterial plasmids carrying AMR determinants.

## Introduction

Bacterial infectious diseases, despite the availability of antibiotics, remain an important public health issue, representing the second leading cause of death worldwide [1]. The gradual increase in the resistance rates of several important bacterial pathogens represents a serious threat to public health [2–4]. Indeed, multidrug-resistant bacteria are the cause of a slow-growing pandemic. The dissemination of multiple antimicrobial resistance (AMR) genes has been largely attributed to the acquisition of plasmids by horizontal gene transfer (HGT), especially in Gram-negative bacteria [5–7], as well as in Gram-positive bacteria [8]. Plasmids can confer resistance to the major classes of antimicrobials [9].

For decades, plasmid incompatibility [10] has been a useful tool for grouping bacterial plasmids. In recent years, other approaches have been considered. Based on phylogenetic analysis of conjugative relaxase, the protein required to initiate plasmid mobilization through conjugation, plasmids can be grouped into different relaxase families [11]. Plasmids belonging to the incompatibility group (Inc) HI (MOB_H_ relaxase family) are widespread in *Enterobacteriaceae* and most commonly include genetic elements encoding multiple AMR determinants [12]. IncHI plasmids, often >200 kb in size, share a common core of approximately 160 kb. The differences in size are due to the presence of distinct insertion elements, including many AMR determinants [13]. IncHI-encoded AMR can be present in enterobacterial such as *Salmonella, Escherichia coli* [14], *Klebsiella pneumoniae* [15] and *Citrobacter freundii* [16]. Plasmids of the IncHI2 subgroup predominate in antibiotic-resistant *Salmonella* isolates. In *S*. Typhi, more than 40% of isolates harbor an IncHI plasmid [17]. In recent years, a novel role of IncHI plasmids in AMR spread has been reported. The emergence of Gram-negative bacteria with AMR, especially those producing carbapenemases, led to reintroduction of colistin as a last resort antibiotic for the treatment of severe infections [18]. In contrast to its limited clinical use, colistin is widely used in veterinary medicine [19]. In the past, colistin resistance was associated with chromosomal mutations only [20]. Nevertheless, plasmid-mediated resistance, conferred by the mobilized colistin resistance gene (*mcr-1*), has emerged recently. Since its discovery in 2016 in China [21], *mcr* genes, have been detected in animals, food, the human microbiota, and clinical samples in over thirty countries [22–26]. IncHI2 plasmids represent 20.5% of the overall plasmids encoding the *mcr-1* gene worldwide but up to 41% in Europe [27]. Of special concern is the presence of the *mcr-1* resistance determinant in *Enterobacteriaceae* carrying carbapenem resistance genes, such as *bla*_*NDM*_ and *bla*_*KPC*_. The combination of these AMR determinants seriously compromises the treatment of infections caused by pathogenic strains harboring these plasmids [28,29]. An example of this is the recent report of an AMR clone of the highly virulent *E. coli* ST95 lineage [14]. *E. coli* ST95 clones underlie neonatal meningitis and sepsis. They are usually sensitive to several antibiotics. This clone harbors an IncHI2 plasmid that encodes, among other factors, determinants of resistance to colistin and multiple other antibiotics (including the extended-spectrum beta-lactamase *blaCTX-M-1*). The spread of such an AMR ST95 clone could pose a threat to human health worldwide [14].

IncHI plasmid conjugation has a distinctive feature: while optimal conjugation rates are obtained at temperatures found outside the host (30°C and below), conjugative transfer is repressed at temperatures encountered within the host (37°C) [30,31]. The plasmid R27 is the prototype of IncHI1 plasmids. It harbors the Tn*10* transposon, which confers resistance to Tc, and has been exhaustively studied. The R27 replication and conjugation determinants are well characterized [32,33], and its complete nucleotide sequence is available [34]. Several ORFs from the plasmid R27 (66%) do not show similarity to any known ORFs.

IncA/C plasmids belong to the same MOB_H_ relaxase family as IncHI plasmids. They were originally identified in the 1970s among multidrug-resistant *Aeromonas hydrophila* and *Vibrio* spp. Isolates that infected cultured fish [35,36]. Since the 1990s, these plasmids have received increasing interest because of their role in mobilizing AMR in enterobacteria and other Gram-negative microorganisms [37–40]. They have an extremely broad host range that includes members of Beta-, Gamma- and Deltaproteobacteria [41] and play a relevant role in the global spread of AMR [42,43]. They represented 50% of all plasmids isolated from *bla*_NDM_-producing *Klebsiella pneumoniae* of clinical origin characterized in a recent study [44].

Proteins containing an immunoglobulin (Ig)-like domain contain several chains of approximately 70-100 amino acid residues present in antiparallel β-strands and organized in two β-sheets that are packed against each other in a β-sandwich. The Ig-like domain has been identified in a large number of proteins with diverse biological functions, is widely distributed in nature, and is present in vertebrates, invertebrates, plants, fungi, parasites, bacteria, and viruses [45]. Bacterial proteins containing Ig-like domains (Big) exhibit a wide range of functions. They include fimbrial subunits, adhesins, membrane transporters and several enzymes (as reviewed in [46]). In a previous report, we studied a high-molecular-weight extracellular protein (the RSP protein) that contains a Big domain and plays an essential role in IncHI plasmids conjugation. Among other targets, the RSP protein appears to be associated with flagella, reducing cell motility. Under specific mating conditions, it could be shown that binding of the RSP protein to the flagella influences conjugation [47]. In this report, we present novel data about the roles of these plasmid-encoded Big proteins. We show that two Big proteins bind both flagella and the conjugative pilus to favor conjugation of the IncHI1 plasmid R27. Furthermore, we also show that other groups of plasmids such as IncA/C and IncP2 also encode these proteins. We provide evidence for their role in the conjugation of IncA/C plasmids. The role of plasmid-encoded Big proteins in plasmid conjugation is discussed.

## Results

### The RSP protein interacts in vitro both with a new R27-encoded Big protein and with a protein involved in plasmid conjugation

To gain further insight into the role of the RSP protein in the conjugation of the R27 plasmid, we decided to assess whether this protein interacts with other proteins expressed by the *Salmonella* strain SL1344 (R27). We performed immunoprecipitation of a cellular extract of strains SL1344 (R27 RSP-Flag) and SL1344 (R27 Δ*rsp*) and analyzed the proteins that specifically coprecipitated with the RSP protein. Two R27-encoded proteins were found to specifically coprecipitate with the RSP protein (Table S1). The protein showing the highest score (187.77) and coverage (58.56) was the R27_p055 protein. The *R0055* gene was mapped between transfer regions 2 and 1 of the R27 plasmid (Fig. 1A). The *R0055* gene product is a 794 AA protein with a molecular mass of 86.75 kDa. As with the RSP protein, the protein encoded by *R0055* also contains bacterial Ig-like domains (Big_1 and _3): a Big_1 domain spanning amino acid residues 143 to 254, and a Big_3 domain spanning residues 537 to 693. The R27_p055 protein, herein termed RSP2, also contains a DUF4165 domain of unknown function (amino acids 23 to 142) (Fig. 1B).

**Figure 1.**
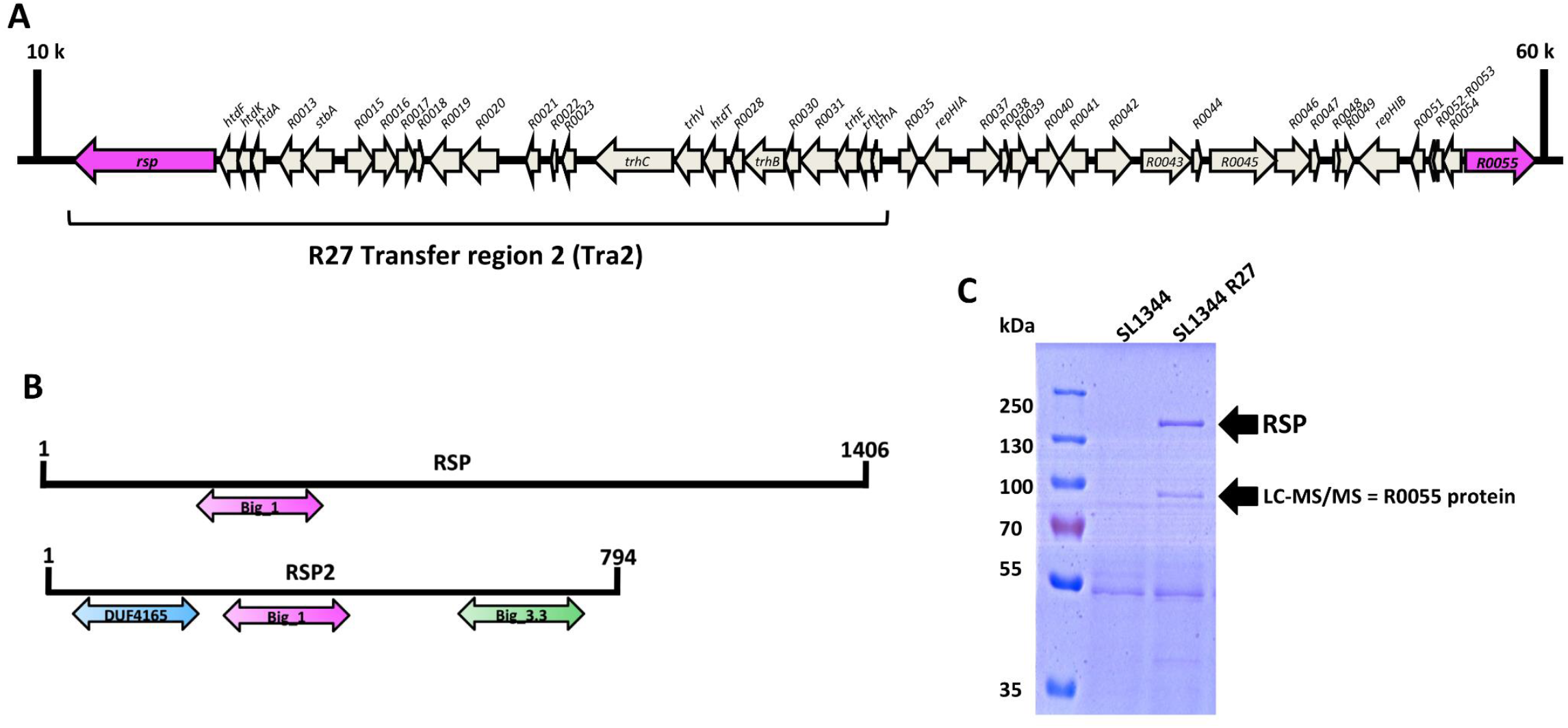
Identification of the RSP2 protein. (A) Genetic map of the R27 plasmid region where both the *rsp* and *rsp2* (*R0055*) genes were mapped. (B) 675 Comparison of the Big domains of the RSP and RSP2 proteins. (C) Detection of the RSP2 protein in the cell-free secretome of the SL1344 (R27) strain. Arrows point to the bands corresponding to the RSP and RSP2 proteins, the latter of which was confirmed by LC-MS/MS analysis.

The second protein showing a high score (59.46) and coverage (39.92) was the R27-encoded TrhH protein. This protein shares 26% of identity with the IncF TraH protein, which is involved in plasmid conjugation [32].

### The RSP2 protein shows RSP-dependent expression in the external medium

To determine whether the RSP2 protein is also present in the cell-free secretome of strain SL1344 (R27), we analyzed the cell-free secreted protein profile of this strain by SDS-PAGE (Fig. 1C). In addition to the band corresponding to the already characterized RSP protein, a second band corresponding to a protein of molecular mass equivalent to that of the RSP2 protein, was observed. The band was isolated and analyzed by LC-MS/MS. It corresponded to the RSP2 protein (Fig. 1C). To identify the RSP2 protein in the different cellular compartments, a Flag-tag was added to the *rsp2* gene (see Materials and Methods section for details). Cultures of strains SL1344 wt and SL1344 (R27 RSP2-Flag) were grown in LB medium at 25°C to an OD_600nm_ of 2.0. Samples were then collected, and the different cellular fractions were obtained. The RSP2 protein was detected in the different fractions by Western blotting, using anti-Flag-specific antibodies (Fig. 2A-C). The protein was identified in the same cell compartments as the RSP protein (i.-e., periplasm, inner membrane, cytoplasm, and cell-free secreted proteins).

**Figure 2.**
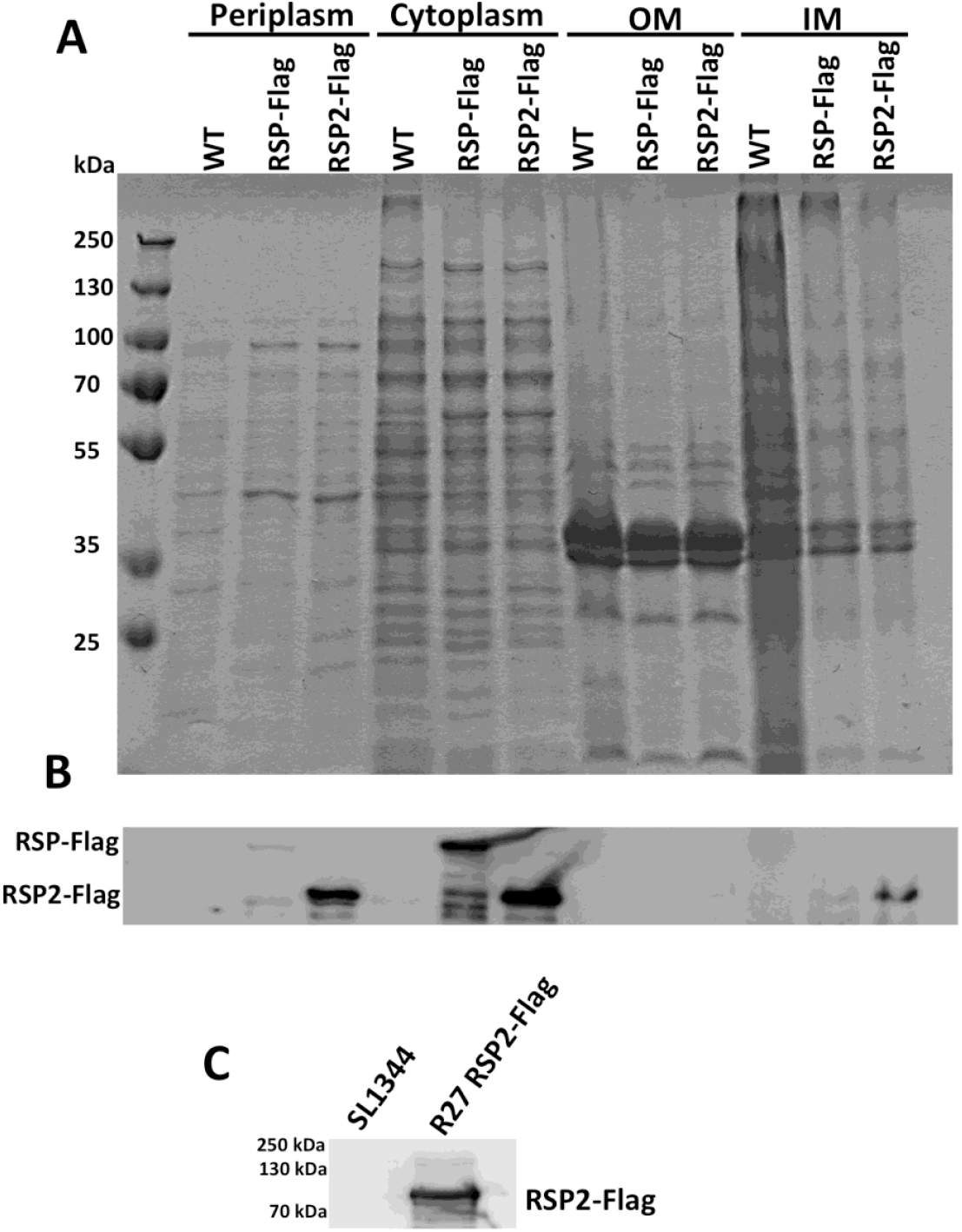
Immunodetection of the RSP2-Flag protein in different cellular compartments. (A) 683 Coomassie blue staining of the different cellular fractions obtained from strains SL11344 wt, 684 SL1344 (R27 RSP-Flag) and SL1344 (R27 RSP2-Flag). (B) Immunodetection of the RSP-Flag and 685 RSP2-Flag proteins in the periplasm, cytoplasm, and outer and inner membrane fractions of 686 strains SL1344 (R27 RSP-Flag) and SL1344 (R27 RSP2-Flag), respectively. (C) Immunodetection of 687 the RSP2-Flag protein in the cell-free secretome of strain SL1344 (R27 RSP2-Flag) grown at 25ºC 688 until OD_600 nm_ of 2.0.

As the above reported data show that the RSP2 protein can be exported to the external medium, we decided to study whether this protein is exported by the R27-encoded type IV secretion system that is also used by RSP [47]. To address this point, we first used the SSPred program (http://www.bioinformatics.org/sspred/html/sspred.html) for *in silico* prediction of whether the RSP2 protein, similar to the RSP protein, could be exported through a type IV secretion system (Fig S1). To provide evidence supporting this hypothesis, we used strains SL1344 (R27 RSP2-Flag) and SL1344 (R27 Δ*trhC* RSP2-Flag) and analyzed the presence of RSP2 protein in the protein profile of the cell-free secreted fractions (secretome). This protein could not be detected in the secretome of strain SL1344 (R27 Δ*trhC*) (Fig. 3A). Immunodetection of RSP2-Flag by using anti-Flag specific antibodies confirmed the requirement for TrhC expression for RSP2 export (Fig. 3B). We next checked whether RSP2 export in strain SL1344 (R27 Δ*trhC*) could be made to occur by providing the gene encoding the TrhC ATPase *in trans*, cloned in the plasmid pBR322 (plasmid pBR322-*trhC*). Complementation of RSP2 export was observed (Fig. 3B), suggesting that, as was the case for the RSP protein, the R27-encoded type IV secretion system mediates export of the RSP2 protein.

**Figure 3.**
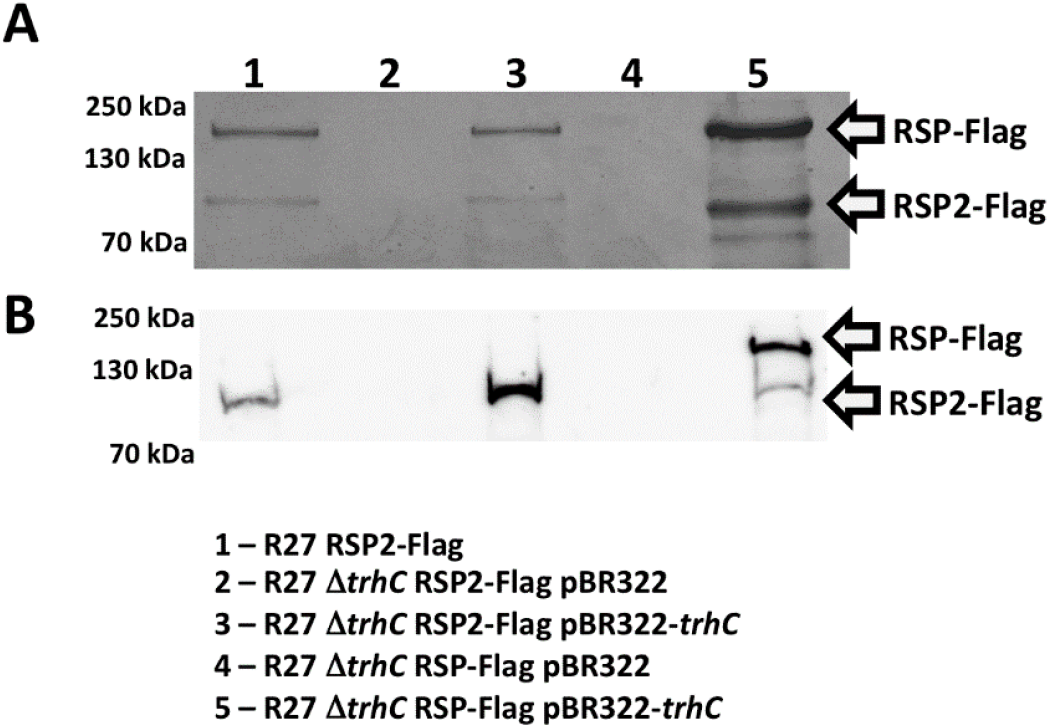
RSP2 export requires the type IV secretion system encoded by the R27 plasmid. (A) Coomassie blue staining for SDS-PAGE analysis of the cell-free secretome of the strains SL1344 (R27 RSP2-Flag), SL1344 (R27 Δ*trhC* RSP2-Flag pBR322), SL1344 (R27 Δ*trhC* RSP2-Flag pBR322-*trhC*), (R27 Δ*trhC* RSP-Flag pBR322) and (R27 Δ*trhC* RSP-Flag pBR322-*trhC*) grown at 25ºC until OD_600 nm_ of 2.0. (B) Immunodetection of the RSP-Flag and RSP2-Flag proteins with anti-Flag antibodies. Arrows point to the RSP-Flag and RSP2-Flag proteins. The experiment was repeated three times. A representative experiment is shown.

We next addressed the question whether there is co-dependence in protein export between the RSP and the RSP2 proteins. To that end, a Flag-tag was added to the *rsp2* gene in the strain SL1344 (R27 Δ*rsp*). Expression of both RSP and RSP2 proteins was assessed in the strains SL1344 (R27 RSP2-Flag), SL1344 (R27 Δ*rsp* RSP2-Flag) and SL1344 (R27 Δ*rsp* RSP2-Flag pLG338-rsp). The results obtained show that export of the RSP2 protein is dependent on the RSP expression (Fig. 4A and 4B).

**Figure 4.**
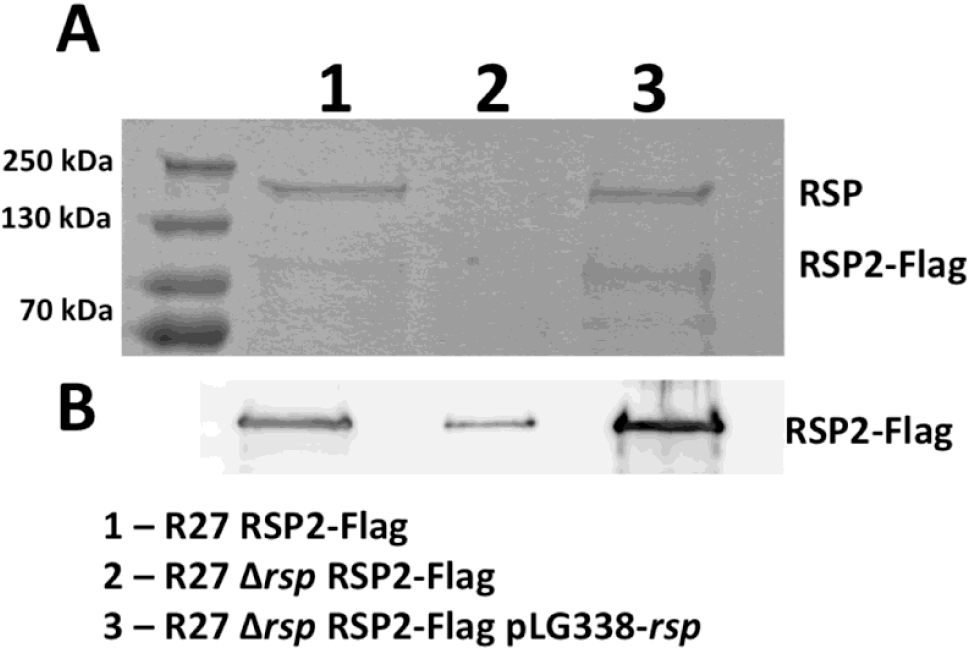
RSP2 export is dependent on the expression of the RSP protein. (A) Coomassie blue 706 staining for SDS-PAGE analysis of the cell-free secretomes of the strains SL1344 (R27 RSP2-Flag), 707 SL1344 (R27 Δ*rsp* RSP2-Flag) and SL1344 (R27 Δ*rsp* RSP2-Flag pLG338-*rsp*) grown at 25ºC until 708 OD_600 nm_ of 2.0. (B) Immunodetection of the RSP2-Flag protein with anti-Flag antibodies. RSP2-709 Flag protein is indicated. The experiment was repeated three times. A representative 710 experiment is shown.

### Expression of the RSP2 protein influences the motility and conjugation of strain SL1344 (R27)

We previously showed that the expression of the RSP protein is essential for R27 plasmid conjugation, and that SL1344 cells that express RSP show reduced motility compared to plasmid-free cells [47]. Considering the observed interaction of both RSP and RSP2 proteins, we studied whether, as is the case for the RSP protein, expression of the RSP2 protein influences cell motility and/or conjugation.

After constructing an R27 derivative lacking the *rsp2* gene (plasmid R27 Δ*rsp2*), we performed a comparative motility assay with the *Salmonella* strain SL1344 and its derivatives harboring the R27, R27 Δ*rsp2*, and R27 Δ*rsp2* pLG338-*rsp2* plasmids. The results obtained (Fig.5A) showed that the RSP2 protein influences the motility of strain SL1344 (R27).

We were able to show previously that the RSP protein is associated with the flagella synthesized by the Salmonella strain SL1344 [47]. We decided next to assess whether RSP2 also targets the flagella. In an attempt to detect the RSP2 protein by transmission electron microscopy, we used strain SL1344 (R27 RSP2-Flag) and gold-labelled anti-Flag monoclonal antibodies (Fig 5B and 5C). As was the case for RSP, RSP2 also exhibited binding to the flagella.

**Figure 5.**
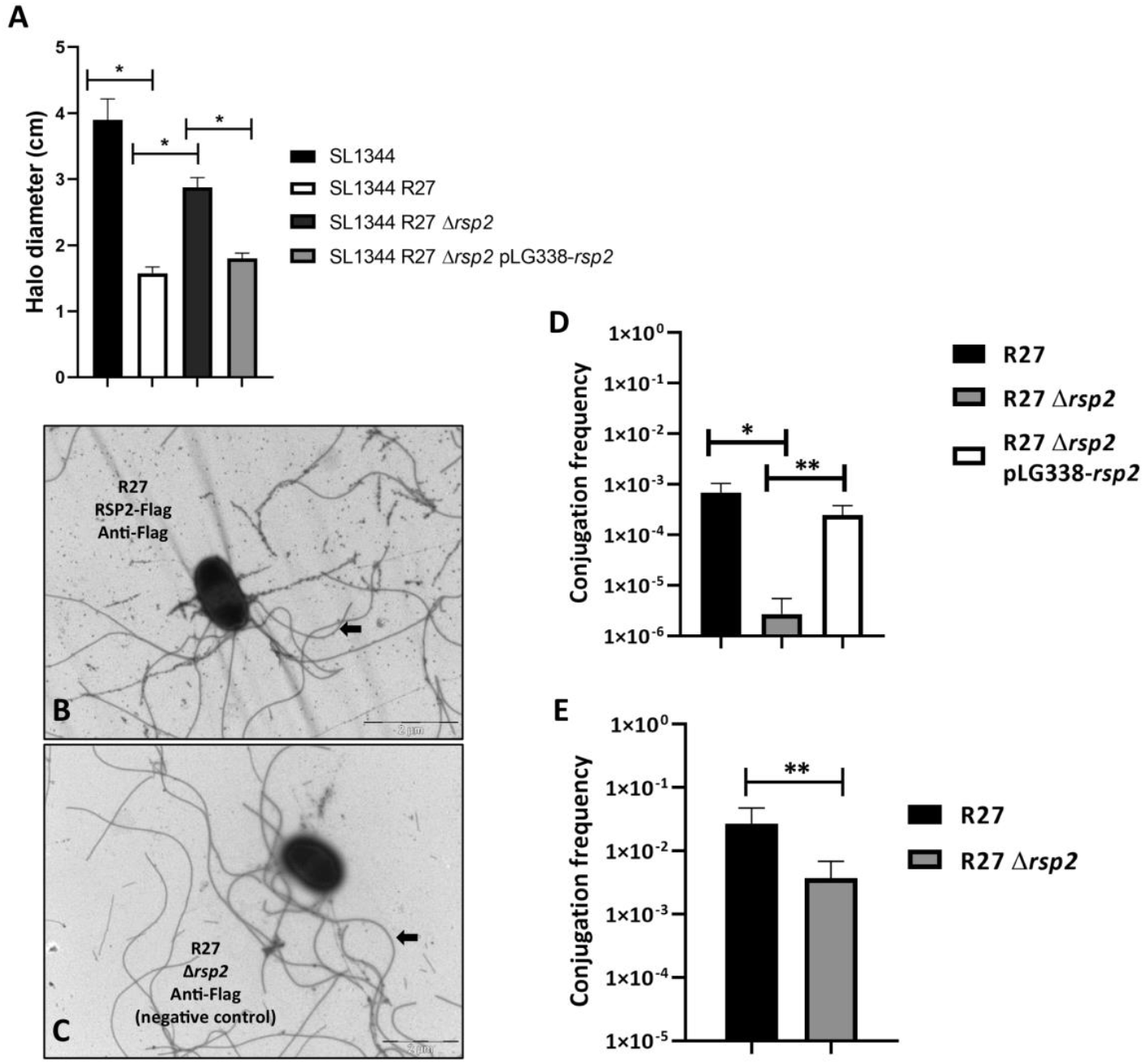
The RSP2 protein influences the motility of strain SL1344 (R27) and the conjugation of 716 the R27 plasmid. (A) Effect of the *rsp2* allele on the motility of strain SL1344 (R27). The results 717 are the means of three independent experiments. Standard deviations are shown. Statistical 718 analysis showed significant differences, analyzed by unpaired two-sided Student’s t-test (\**P*-719 value < 0.0001). (B and C) The RSP2 protein binds to the flagella of strain SL1344 (R27). 720 Immunogold electron microscopy of cells from strains SL1344 (R27 RSP2-Flag) (B) and SL1344 721 (R27 Δ*rsp2*) (C) using monoclonal anti-Flag antibodies and goat anti-mouse IgG conjugated to 12 722 nm gold particles. Arrows point to the RSP protein associated with the flagella in (B) and only to 723 the flagella in (C). Bars represent 2 μm. (D and E) Effect of the *rsp2* allele on the conjugation 724 frequency of the R27 plasmid in liquid (D) and solid (E) media, respectively. The data shown are 725 the means and standard deviations of three independent experiments. Statistical analysis 726 showed significant differences, analyzed by unpaired two-sided Student’s t-test (D) (\**P*-value 0.0082, \*\**P*-value 0.0095) and (E) (\*\**P*-value 0.0428).

To assess the role of the RSP2 protein on plasmid conjugation, we constructed a R27 derivative lacking the *rsp2* gene (plasmid R27 Δ*rsp2*) and compared the conjugation frequencies of strains SL1344 (R27), SL1344 (R27 Δ*rsp2*) and SL1344 (R27 Δ*rsp2* pLG338-*rsp2*) growing at 25°C in liquid media. We also compared the conjugation frequencies of strains SL1344 (R27) and SL1344 (R27 Δ*rsp2*) growing cells on nitrocellulose filters placed on LB plates. When cells were grown in liquid medium, transfer of the R27 Δ*rsp2* plasmid was detected at a frequency that was approximately two logs lower than that of wt R27 (Fig.5D). The presence *in trans* of the RSP2 protein encoded by the pLG338-*rsp2* plasmid, restored the conjugation frequency of the wt R27 plasmid. When cells were grown on solid medium, we also detected a significant decrease in the conjugation frequency of the R27 Δ*rsp2* plasmid compared to that of the wt R27 plasmid (Fig. 5E).

### Relationship between the RSP and RSP2 proteins and the conjugative machinery of the R27 plasmid

As mentioned above, immunoprecipitation of the RSP protein indicated interactions with both the RSP protein and the R27 *trhH* gene product. The TrhH protein of the plasmid R27 shares identity with the TraH protein encoded by IncF plasmids [32]. The TraH protein is a component of the outer membrane complex involved in conjugation [48] and has been shown to be required for pilus assembly [49]. On the other hand, the *trhA* gene product encoded in IncHI plasmids has been considered to be the pilin subunit itself [50]. We decided to analyze whether the expression of the R27 TrhH and/or TrhA proteins encoded by IncHI plasmids influences the expression of the RSP and RSP2 proteins. After inactivation of the *trhH* or *trhA* genes of strain SL1344 (R27), the expression of the RSP and RSP2 proteins in the external medium was analyzed. As both the RSP and RSP2 proteins were copurified with flagella when a conventional flagella purification protocol was used (see the Material and Methods section), we obtained fractions containing flagella from the SL1344 wt, SL1344 (R27), SL1344 (R27 Δ*trhA*) and SL1344 (R27 Δ*trhH*) isogenic derivatives. Proteins were analyzed by SDS-PAGE (Fig. 6A). The expression of both the RSP and RSP2 proteins in the extracellular medium is dependent upon the function of TrhH or TrhA. We also analyzed the intracellular expression of the RSP and RSP2 proteins in the *trhH* and *trhA* genetic backgrounds. Lack of either TrhH (Fig. 6B and 6C) or TrhA (Fig. 6D and 6E) function resulted in intracellular accumulation of either RSP or RSP2. Considering that expression of the RSP and RSP2 proteins reduces cell motility in strain SL1344 (R27) and that inactivation of the *trhH* gene interferes with the presence of these proteins in the extracellular medium, it could be expected that *trhH* inactivation would also result in increased motility in SL1344 cells harboring plasmid R27 Δ*trhH*. We then performed a mobility assay of the plasmid-free SL1344 strain and of its SL1344 (R27) SL1344 (R27 Δ*rsp*), SL1344 (R27 Δ*rsp2*) and SL1344 (R27 Δ*trhH*) derivatives. In accordance with (i) the observed requirement of TrhH function for RSP and RSP2 export and (ii) the effect of the RSP/RSP2 proteins on cell motility, the motility of strain SL1344 (R27 Δ*trhH*) was significantly reduced compared to that of the wt strain (Fig. 7).

**Figure 6.**
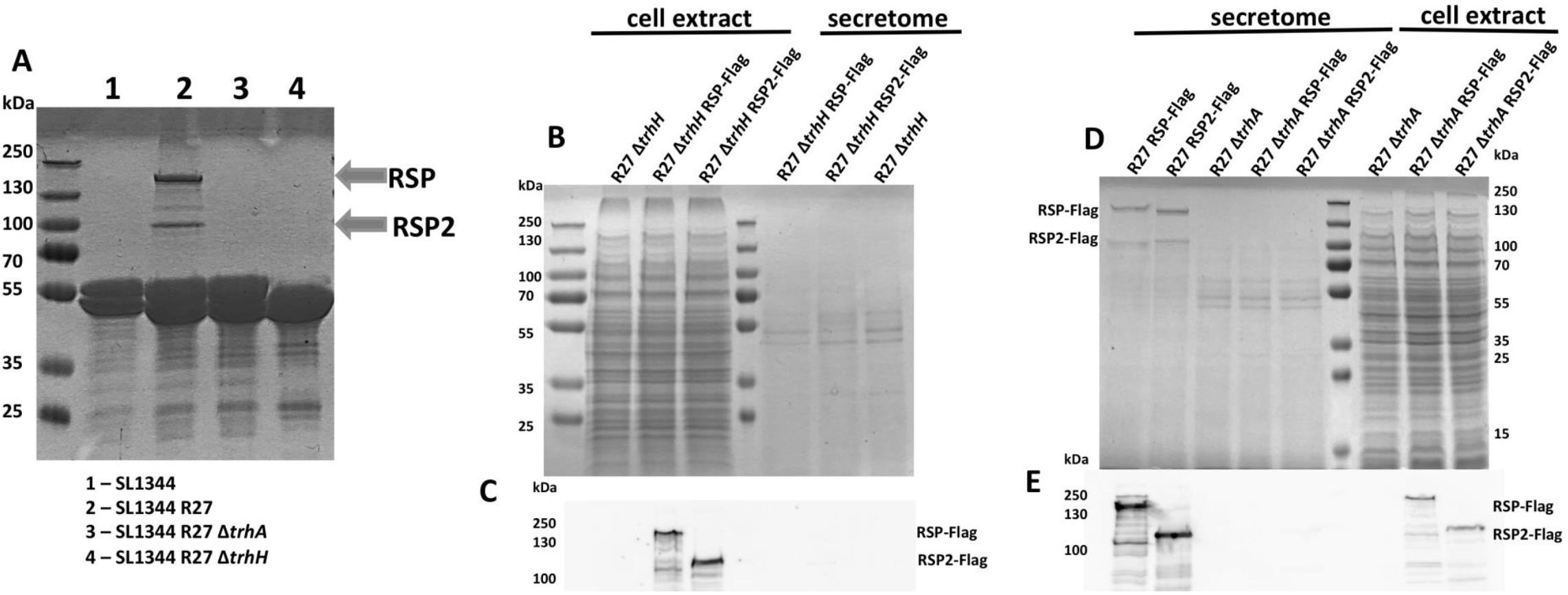
Expression of the RSP and RSP2 proteins depends on the R27 TrhA and TrhH functions. (A) SDS-PAGE analysis of the purified flagellar fractions of the plasmid-free strain SL1344 and strain SL1344 harboring (R27) and the (R27 Δ*trhA*) and (R27 Δ*trhH*) derivatives. (B) SDS-PAGE analysis of the cell extract and cell-free secretome of the strains SL1344 (R27 Δ*trhH*), SL1344 (R27 Δ*trhH* RSP-Flag) and SL1344 (R27 Δ*trhH* RSP2-Flag), respectively. (C) Immunodetection of RSP-Flag and RSP2-Flag in the intracellular compartments of strain SL1344 harboring the corresponding *trhH* derivatives of the R27 plasmid. (D) SDS-PAGE analysis of the cell extract and cell-free secretome of the strains SL1344 (R27 RSP-Flag), SL1344 (R27 RSP2-Flag), SL1344 (R27 Δ*trhA*), SL1344 (R27 Δ*trhA* RSP-Flag) and SL1344 (R27 Δ*trhA* RSP2-Flag), respectively. (E) Immunodetection of RSP-Flag and RSP2-Flag in the intracellular compartments of strain SL1344 735 harboring the corresponding *trhA* derivatives of the R27 plasmid. The experiments were repeated three times. A representative experiment is shown.

**Figure 7.**
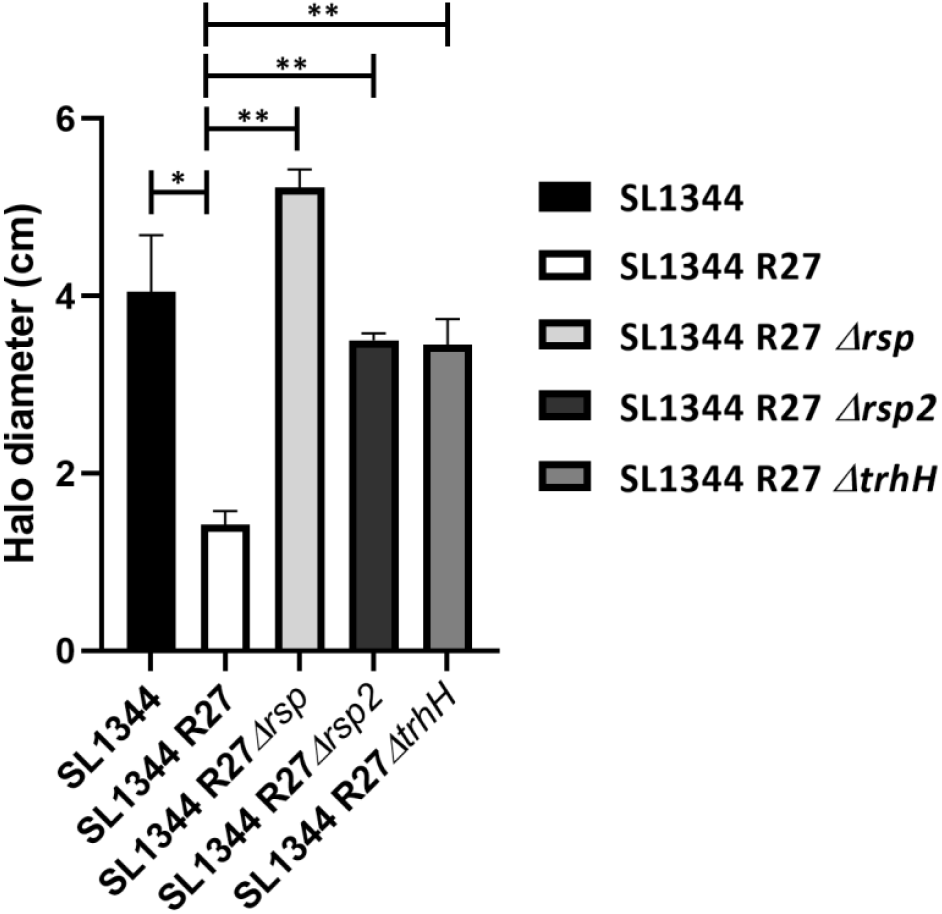
Loss of TrhH function impairs the effect of the RSP and RSP2 proteins on SL1344 cell 742 motility. The motility of the different strains was measured as the halo diameter of the different 743 colonies growing on motility agar. The results are the means of three independent experiments. 744 Standard deviations are shown. Statistical analysis showed a significant difference analyzed by 745 unpaired two-sided Student’s t-test (\**P*-value 0.0009, \*\**P*-value <0.0001).

### The RSP and RSP2 proteins bind the R27 conjugative pili

Taking into account the observed relationship between elements of the conjugation machinery of the R27 plasmid and the RSP and RSP2 proteins, we decided to assess whether these proteins, in addition of binding the flagella, they also target the conjugative pilus. To that end, we blocked flagella expression and used electron microscopy on order to detect either the RSP or RSP2 proteins bound to the conjugative pili. To prevent flagella expression, we constructed Δ*fliC/fljB* (flagellin subunit) derivatives of strains SL1344 (R27) and SL1344 (R27 RSP2-Flag). Immunogold transmission electron microscopy imaging of these strains by using either polyclonal anti-RSP antibodies [47] or monoclonal anti-Flag antibodies showed that in both examples, gold particles were associated with tubular structures that likely corresponded to the conjugative pilus (Fig. 8A and 8C). When the Δ*trhH* allele was introduced into the mutant derivatives lacking flagella, the tubular structures were no longer detected, thus supporting the hypothesis that they corresponded to the conjugative pilus (Fig. 8B and 8D).

**Figure 8.**
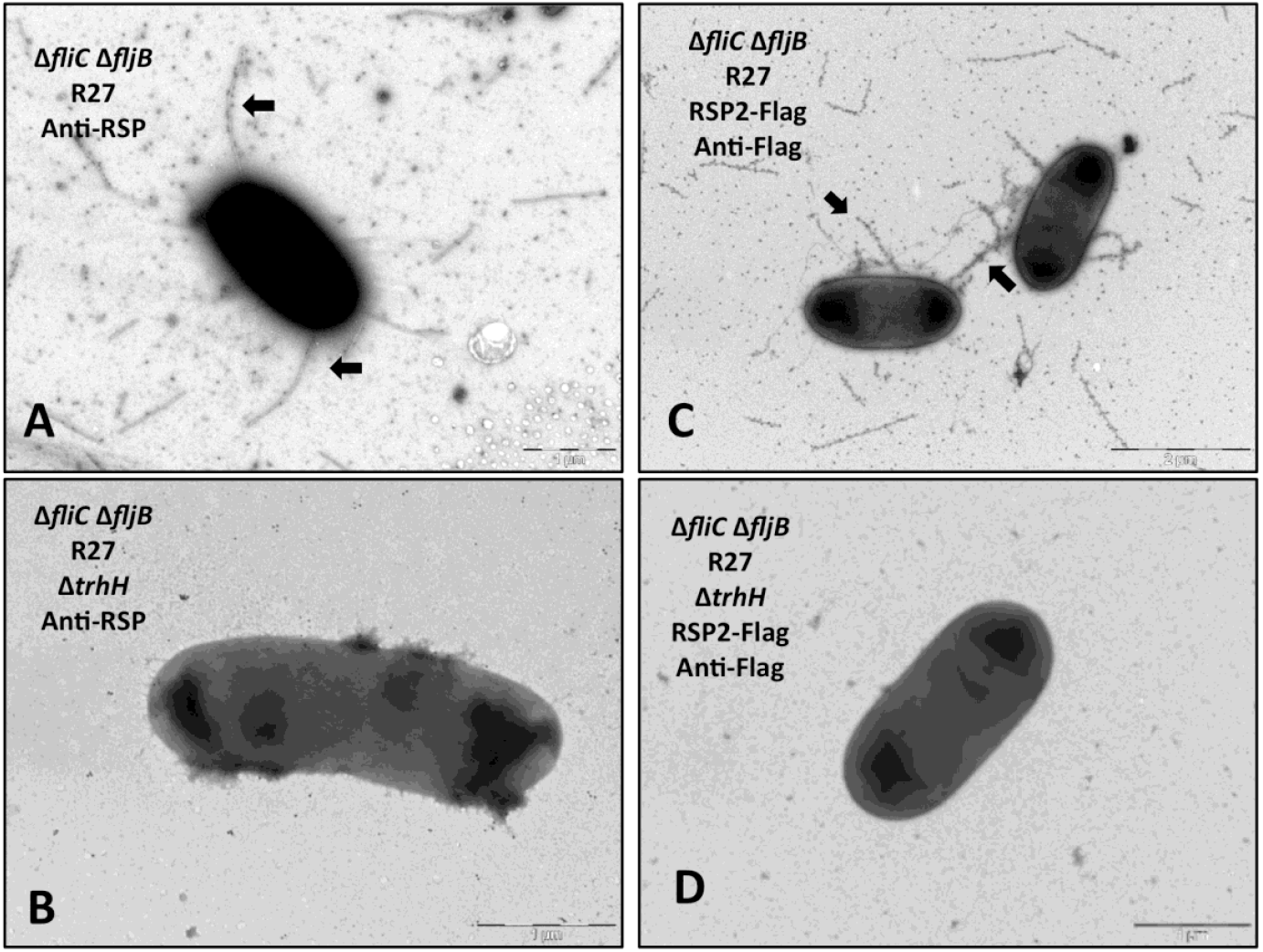
The RSP and RSP2 proteins bind to the conjugative pilus encoded by the R27 plasmid. 751 Immunogold electron microscopy of cells from strains SL1344 Δ*fliC* Δ*fljB* (R27) (A) and SL1344 752 Δ*fliC* Δ*fljB* (R27 Δ*trhH*) (B) using polyclonal anti-RSP antibodies and goat anti-rabbit IgG 753 conjugated to 12 nm gold particles. Arrows point to the RSP protein associated with the 754 conjugative pilus. Immunogold electron microscopy of cells from strains SL1344 Δ*fliC* Δ*fljB* (R27 755 RSP2-Flag) (C) and SL1344 Δ*fliC* Δ*fljB* (R27 RSP2-Flag Δ*trhH*) (D) using anti-Flag monoclonal 756 antibodies and goat anti-mouse IgG conjugated to 12 nm gold particles. Arrows point to the 757 RSP2 protein associated with the conjugative pilus. Bars represent 1 μm (in A, B and D) and 2 758 μm (in C).

### The RSP2 protein is specific for IncHI1 plasmids

The RSP protein is restricted to the IncHI plasmids, from both the IncHI1 and IncHI2 subgroups [47]. Upon having shown the relevant role of the RSP2 protein in the conjugation of the IncHI1 plasmid R27, we performed a BLAST search to identify the *rsp2* gene in other bacterial plasmids (Table S2). In contrast to the RSP protein, the RSP2 protein is present in IncHI1 plasmids but not in IncHI2 plasmids. Interestingly, a group of IncN plasmids also contains an homolog of the R27 *rsp2* gene.

### Distribution of proteins containing the Big domain among bacterial plasmids

We next addressed the question of whether proteins containing Big domains are a feature of only IncHI plasmids or whether they are also encoded by plasmids of other incompatibility groups. We used for this analysis the genome viewer integrated into the NCBI database, configuring it to show the features and domains of the annotated proteins. The search was performed in assembled and sequenced plasmids of all incompatibility groups, selecting those that presented proteins with annotated Big domains. Among all the plasmids analyzed, proteins with these domains were also found in the plasmids of the IncA/C and IncP2 incompatibility groups. Big proteins from IncA/C plasmids show a high degree of similarity (Fig. S2) and can be mapped to the corresponding *tra* regions (Fig. S3 and Fig. S4). Big proteins from IncP2 plasmids also show a very high degree of similarity (Fig. S5) and can be mapped close to a pilin gene (Fig. S4).

### The ALG87338.1 gene product of the IncA/C plasmid pKAZ3 is required for plasmid conjugation at low temperature

Upon having identified genes encoding proteins containing Big domains harbored on plasmids different from those of the IncHI group, we also aimed to assess whether a protein containing a Big domain encoded by a plasmid different from the IncHI group is also found in the secretome and influences conjugation. For this study, we selected the IncA/C plasmid pKAZ3, encoding the ALG87338.1 gene, the product of which contains a Big domain. The plasmid pKAZ3 was isolated from an antibiotic-contaminated lake and confers multiple-antibiotic resistance [51]. The ALG87338.1 gene product is an 1843 AA protein (200.75 kDa) that contains Big (3_2 and 3_3) and DUF4165 domains (Fig. S6). Plasmid pKAZ3 was first transferred to strain SL1344, and the secretomes of strains SL1344 and SL1344 (pKAZ3) were compared. A band corresponding to a protein of a molecular mass corresponding to the ALG87388.1 gene product was identified (Fig. 9A). This was confirmed by LC-MS/MS analysis of the protein excised from the SDS-PAGE gel. We next generated an ALG87388.1 knockout mutant derivative of plasmid pKAZ3. Considering that expression of either the RSP or RSP2 protein reduces SL1344 cell motility, we also compared the motility of strains SL1344, SL1344 (pKAZ3) and SL1344 (pKAZ3 ΔALG87388.1) (Fig. 9B). We observed that, while the acquisition of the pKAZ3 plasmid reduced cellular motility, there were no significant differences in motility between strains SL1344 (pKAZ3) and SL1344 (pKAZ3 ΔALG87388.1). We also compared the conjugation frequencies of strains SL1344 (pKAZ3) and SL1344 (pKAZ3 ΔALG87388.1), at both 37°C and at 25°C (Fig. 9C). ΔALG87388.1 did not influence the conjugation frequency at 37°C, but it had a very strong effect at 25°C.

**Figure 9.**
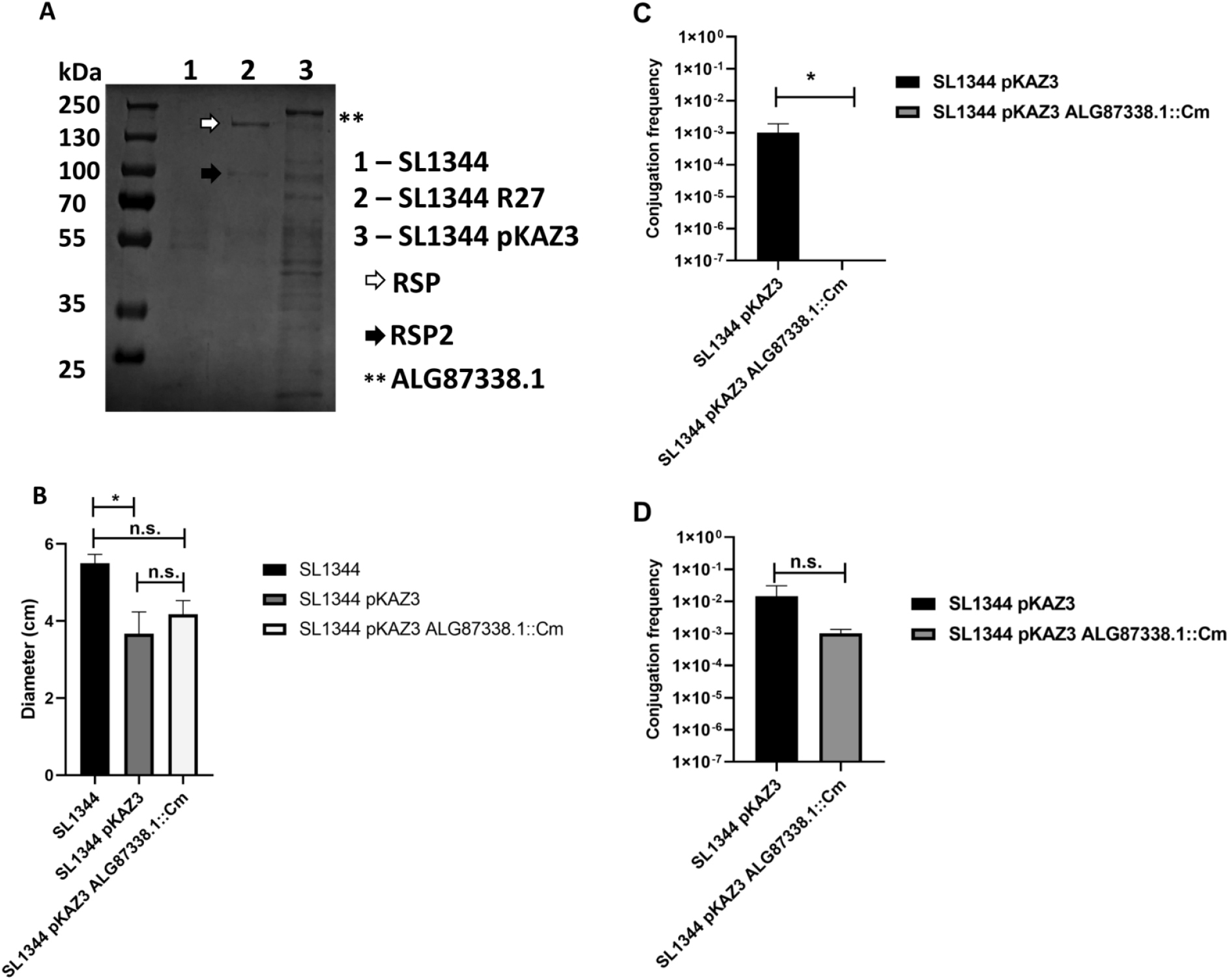
The ALG87388.1 gene product of the pKAZ3 plasmid influences conjugation at 25°C. (A) SDS-PAGE analysis of the cell-free secretomes of strains 762 SL1344, SL1344 (R27) and SL1344 (pKAZ3) grown at 25ºC until OD_600 nm_ of 2.0. Asterisks indicated the detection of the ALG87388.1 protein in the secretome 763 of strain SL1344 (pKAZ3) confirmed by LC-MS/MS. Arrows indicate the RSP and RSP2 protein. (B) Motility of the SL1344, SL1344 (pKAZ3) and SL1344 (pKAZ3 764 ΔALG87388.1) strains. The data shown are the means and standard deviations of three independent experiments. Statistical analysis showed significant 765 differences analyzed by unpaired two-sided Student’s t-test (\**P*-value 0.0469; n.s. not significant). (C and D). Effect of the ΔALG87388.1 allele on the 766 conjugation frequency of the pKAZ3 plasmid at 25°C (C) and 37°C (D) in liquid media. The data shown are the means and standard deviations of three 767 independent experiments. Statistical analysis showed significant differences analyzed by unpaired two-sided Student’s t-test (\**P*-value 0.042; n.s. not 768 significant).

## Discussion

The conjugative process has been studied for decades, and a detailed picture of the molecular mechanism of conjugational DNA transfer is available (as reviewed in [50,52,53]). Plasmids such as the F factor, RP4, R388 or pTi have been studied as reference models to determine the function of several Tra proteins in plasmid conjugation by using genetic and biochemical approaches [50]. Comprehensive information is available on processes such as conjugative pilus biosynthesis, the establishment of donor-recipient cell contact or the assembly and activity of the relaxosome, which are key aspects of bacterial conjugation that must occur prior to the DNA transfer process. Nevertheless, several key questions, such as those regarding the function of several Tra proteins or the pilus’s ability to transport DNA between distant donor and recipient cells, remain to be answered [50]. In this report, we have elaborated on the role of plasmid-encoded high-molecular-weight Big proteins in the conjugation process of different plasmids.

In the IncHI plasmid R27, two genes that were mapped to the Tra2 and Tra1-Tra2 intergenic regions (*rsp* and *rsp2*, respectively) [32,33], encode proteins containing Big domains that participate in the conjugation process. Both the RSP and RSP2 proteins bind flagella, this resulting in a reduced cell motility. Export of the RSP2 protein to the external medium appears to be dependent upon RSP export. This may explain that, whereas loss of RSP function completely compensates for the effect of R27 acquisition on bacterial cell motility, loss of RSP2 function only shows a partial effect on cell motility.

The relationship between the RSP and RSP2 proteins, and the conjugative machinery of the R27 plasmid is shown in this work. Expression of key elements required for the synthesis of the conjugative pilus of incHI1 plasmids (TrhH protein) and of the pilin subunit itself (TrhA protein) is required for the correct translocation of the RSP and RSP2 proteins to the external surface of the cells. In addition, imaging of cells lacking flagella showed short filaments that likely corresponded to the conjugative pilus. These filaments were targeted both by the RSP and RSP2 proteins. Hence, there is an interaction between the RSP and RSP2 proteins and the R27 conjugative pilus.

IncHI plasmids and *Salmonella* participate in regulatory crosstalk that, is based on the expression of both the tetracycline determinant and other plasmid-encoded genes [54,55]. One of the effects of the crosstalk is the alteration in cellular motility, which is mediated by both plasmid-borne regulators [56] and the binding of the RSP and RSP2 proteins to the flagellar structure ([47], this work). Nevertheless, these proteins also play a second and critical role in IncHI plasmid conjugation. They also bind the conjugative pilus, which is likely required to facilitate transmission of the conjugative plasmid. Plasmid-encoded Big proteins binding to the flagella and thus reducing motility may favor cell-to-cell contact and hence conjugation. Binding to the conjugative pilus may contribute to the stabilization of the pilus structure. The overall effect of the expression of these proteins is to significantly favor the transfer of the plasmid that encode them.

Genes encoding proteins containing Big domains are not restricted to IncHI plasmids. The genomic analysis performed in this work has shown that such proteins are also present in IncA/C and IncP2 plasmids. Both groups of plasmids have a wide host range. IncA/C plasmids are also key players in the dissemination of AMR in *Enterobacteriaceae* and other Gram-negative microorganisms. IncP2 plasmids are high-molecular-weight plasmids that are prevalent in *Pseudomonas* and contribute to the dissemination of AMR within this genus [57,58]. As the search that we performed had some limitations, it cannot be ruled out that other groups of plasmids also express Big proteins. By using the genome viewer integrated into the NCBI database, only those plasmids that were well-annotated and characterized were considered. In addition, proteins that contain Big domains were annotated because of the confirmation that this domain, was present, not because they shared amino acid sequence similarity. Notably, the RSP2 protein is specific of a subgroup of IncHI plasmids, incHI1. IncHI2 plasmids that encode the *rsp* gene lack the *rsp2* gene. Interestingly, our BLAST analysis also showed that the *rsp2* gene has jumped to plasmids of a different Inc group, IncN. This may indicate that the *rsp2* genes are spreading and may also influence conjugation in these plasmids.

The identified high-molecular-weight Big proteins encoded by IncA/C plasmids show a very high degree of homology, which suggests that they play similar roles in these plasmids. The genes encoding these proteins were also mapped to the *tra* region *(traV/A/W/F/N* genes) and, in some instances, were adjacent to the pilin genes. A recent study focused on performing a comprehensive analysis of IncC plasmid conjugation [59], and the gene encoding the identified Big protein encoded by the pMS6198A plasmid (the product of the gene MS6198_A094) was considered not to influence conjugation, despite mapping within the Tra1 region, between the *dsbC*A and *traL* genes. The likely reason for this was that the conjugation experiments were performed only at 37°C. We showed here that the ALG87338.1 gene product of the IncA/C plasmid pKAZ3 positively influenced conjugation at 25°C (i.e., at environmental temperature). IncHI and IncA/C plasmids belong to the same relaxase family (MOB_H_). Interestingly, they also share the requirement for a high-molecular-weight Big protein for efficient conjugation at environmental temperatures. The identified Big proteins of IncP2 plasmids also show a high degree of similarity. While the molecular mass of these proteins is lower than that of the Big proteins of IncHI and IncA/C plasmids, they map close to a pilin, which also suggests that they may play a role in the conjugative process.

Out of the IncHI plasmids, which have classically been characterized as being conjugative only at temperatures below 30°C, plasmids harbored by *Enterobacteriaceae* have usually been studied in cells growing at the optimal growth temperature for these microorganisms, that is, 37°C. Our study also highlights the importance of studying plasmid conjugation at other temperatures, closer to those that most microorganisms encounter outside their warm-blooded hosts. Indeed, AMR transmission occurs mainly in natural environments at temperatures that rarely reach 37°C.

The relevance of the Big proteins studied in this work is not only their role as elements favoring plasmid conjugation but also the fact that, as they are associated with extracellular appendages of the bacterial cells, they can be targeted with specific antibodies either to restrict the dissemination of the plasmids that encode them or, when expressed within the human body, to control infections caused by bacteria that express the plasmid that encode these proteins.

## Materials and Methods

### Bacterial strains, plasmids, and growth conditions

The bacterial strains (Table S3) were routinely grown in Luria-Bertani (LB) medium (10 g l^-1^ NaCl, 10 g l^-1^ tryptone and 5 g l^-1^ yeast extract) with vigorous shaking at 200 rpm (Innova 3100, New Brunswick Scientific). The antibiotics used were chloramphenicol (Cm) (25 μg ml^-1^), tetracycline (Tc) (15 μg ml^-1^), carbenicillin (Cb) (100 μg ml^-1^) and kanamycin (Km) (50 μg ml^-1^) (Sigma-Aldrich).

### Oligonucleotides

The oligonucleotides (from 5’ to 3’) used in this work are listed in Table S4.

### Genetic manipulation

All enzymes used to perform standard molecular and genetic procedures were used according to the manufacturer’s recommendations. To introduce plasmids into *E. coli* and *Salmonella*, bacterial cells were grown until an OD_600 nm_ of 0.6. Cells were then washed several times with 10% glycerol, and the respective plasmids or DNA was electroporated by using an Eppendorf gene pulser (Electroporator 2510).

Deletions of the *rsp2* (ORF *R0055), fliC, fljB, trhH* and *trhA* genes were performed in strain SL1344 (R27) by using the λ Red recombination method, as previously described [60]. The antibiotic resistance determinant of the plasmids pKD3/pKD4 was amplified using the corresponding oligonucleotides (P1/P2 series, see Table S4). The mutants were confirmed by PCR using the corresponding oligonucleotides (P1up/P2down series, see Table S4). We used phage P22 HT for combining mutations by transduction [61]. When necessary, the antibiotic resistance cassette was eliminated using the FLP/FRT-mediated site-specific recombination method as previously described [62].

For deletion of the ALG87338.1 gene from the pKAZ3 plasmid, we first constructed the plasmid pKD46-Km^R^ following the strategy described in [63]. Briefly, kanamycin resistance from the pKD4 plasmid was amplified using the PkmXmnIFW/PkmXmnIRv oligonucleotides together with Phusion Hot Start II High-Fidelity DNA Polymerase (Thermo Scientific) following the manufacturer’s recommendations. The corresponding XmnI-flanked kanamycin resistance fragment was cloned into the vector pKD46 previously digested with the same enzyme. The resulting plasmid was termed pKD46-Km^R^. Deletion of ALG87338.1 was performed in strain SL1344 (pKAZ3) pKD46-Km^R^ by using the λ Red recombination method. The antibiotic resistance determinant of the plasmid pKD3 was amplified using the corresponding oligonucleotides (P1/P2 series, see Table S4). The mutants were confirmed by PCR using the corresponding oligonucleotides (P1up/P2down series, see Table S4).

Recombinational transfer of the Flag sequence into the *rsp2* gene was achieved by following a previously described methodology [64]. The template vector encoding Flag and Km^r^ used was the pSUB11 plasmid. The primers used for construction of the Flag-tagged derivative were R27_p0553XP1 and R27_p0553XP2 (Table S4). The correct insertion of the Flag-tag was confirmed by PCR using oligonucleotides R27_p0553XP1UP and R27_p0553XP2DOWN (Table S4).

To construct the plasmid pLG338-*rsp2*, the ORF *R0055* (*rsp2*) (GenBank accession number AF250878.1, position 56639-59970) was amplified using the oligonucleotides R27_R55 pLG322 ECORI fw/R27_R55 pLG322 Bam rv (see Table S4 for the sequences) together with Phusion Hot Start II High-Fidelity DNA Polymerase (Thermo Scientific) following the manufacturer’s recommendations. The corresponding EcoRI/BamHI fragment was cloned into the vector pLG338-30 previously digested with the same enzymes. The resulting plasmids were Sanger sequenced and termed pLG338-*rsp2*.

### Plasmid conjugation

The R27 and pKAZ3 plasmids were conjugated as described previously [47]. The mating frequency was calculated as the number of transconjugants per donor cell. Student’s *t*-test was used to determine statistical significance, and the values were obtained by using GraphPad Prism 8 software. A *P* value of less than 0.05 was considered significant.

### Motility assays

The motility assay was performed as previously described [47]. The experiments were repeated three times with three plates of each strain in each experiment. The colony diameter was measured and plotted, and standard errors were calculated. Student’s t-test was used to determine statistical significance, and the values were obtained by using the GraphPad Prism 8 software. A P value of less than 0.05 was considered significant.

### Flagellum Isolation

Flagellum isolation was prepared as previously described [47].

### Cell-free secreted proteins (secretome)

Cell-free supernatants were prepared as previously described [47].

### Cell fractionation

Cell fractionation was performed as previously described [47].

### Immunogold electron microscopy

Immunogold microscopy experiments were performed as previously described [47].

### Electrophoresis and Western blotting analysis of proteins

Protein samples were analyzed by 10% and 12.5% SDS-PAGE [65]. Proteins were transferred from the gels to PVDF membranes using the Trans-Blot Turbo system (Bio-Rad). Western blot analysis was performed with a monoclonal antibody raised against the Flag-epitope (Sigma) diluted 1:10.000 in a solution containing PBS, 0.2% Triton, and 3% skim milk and incubated for 16 hours at 4°C. The membranes were washed for 20 minutes each with PBS and 0.2% Triton solution. The washing step was repeated three times. Thereafter, the membranes were incubated with horseradish peroxidase-conjugated goat anti-mouse IgG (Invitrogen) diluted 1:5.000 in a solution containing PBS and 0.2% Triton for 1 hour at room temperature. Again, three washing steps of 45 minutes with PBS and 0.2% Triton solution were performed, and detection was performed by enhanced chemiluminescence using ImageQuant LAS 54000 imaging system software (GE Healthcare Lifesciences).

### RSP-Flag Immunoprecipitation

For RSP-Flag protein immunoprecipitation, the strains SL1344 (R27 RSP-Flag) and SL1344 (R27 Δ*rsp*, negative control) were grown in LB medium for 16 hours at 37ºC. One hundred milliliters of fresh LB medium was inoculated 1:100 with both overnight-cultured strains and grown at 25°C until an OD_600 nm_ of 2.0 was reached. Then, the cells were centrifuged at 9.000 rpm for 30 minutes at 4°C, the pellets were discarded, and the supernatants were filtered through 0.22 μm filters. For each immunoprecipitation protocol, we used 100 μl of Anti-Flag M2 Affinity Gel (Sigma-Aldrich) and 100 ml of each supernatant and incubated the mixture under slow rotation at 4°C for 16 hours. Each supernatant was loaded onto a Poly-Prep chromatography column (Bio-Rad), and the flowthrough was stored at −20°C for further analysis. Each column was washed out with 100 ml of washing buffer (50 mM Tris-HCl pH 7.5, 150 mM NaCl), and elution was performed 3 times with 0.3 ml of elution buffer (0.1M glycine pH 3.5). Elution fractions were concentrated using a trichloroacetic acid (TCA) precipitation protocol, briefly, 1 ml of the samples was mixed with 0.5 ml of a 45% TCA solution (w/v). The samples were kept on ice for 30 minutes and centrifugated for 30 minutes at 13,400 rpm. The supernatants were carefully discarded, acetone was added to the protein pellets, and the pellets were again centrifugated for 30 minutes at 13,400 rpm. The supernatants were carefully discarded. The protein pellets were dried and solubilized with 1x Laemmli Sample Buffer (Bio-Rad). Samples were boiled for 10 minutes and loaded onto a 12.5% SDS-PAGE gel. When the protein samples entered the stacking phase, the gel run was stopped, and samples were excised from the gel and sent to the Proteomic Platform (Barcelona Science Park, Barcelona, Spain) for protein identification, as previously described [55]. Immunoprecipitation and protein identification experiments were repeated twice.

## Acknowledgments and funding sources

This work was supported by grants from Fundació “La Marató TV3”, Spain (Project ref. 201818 10), and PID2019-107479RB-I00 (AEI/FEDER, UE) from the Ministerio de Economía, Industria y Competitividad, and CERCA Programme/Generalitat de Catalunya to A.J. Authors wish to thank Dr. Álvaro San Millán (CNB-CSIC, Madrid) who kindly provided the pKAZ3 plasmid.

