## Supplementary Information for "Roles of proteins containing immunoglobulin-like domains in the conjugation of bacterial plasmids"

| | Accession | Description | Score | Coverage | # Proteins | Unique Peptid | # Peptides | # PSMs | # AAs | MW [kDa] | calc. pI |
|---|---|---|---|---|---|---|---|---|---|---|---|
| RSP-Flag | Q9L5V1 | Uncharacterized protein OS=Salmonella typhi OX=90370 GN=R0009 PE=4 SV=1 - [Q9L5V1_SALTI] - **RSP** | 532,34 | 69,77 | 4 | 84 | 84 | 184 | 1406 | 155,4 | 5,47 |
| | A0A0W4T704 | Uncharacterized protein OS=Salmonella enterica OX=28901 GN=IN95_04195 PE=4 SV=1 - [A0A0W4T704_SALER] - **RSP2** | 187,77 | 58,56 | 2 | 35 | 35 | 62 | 794 | 86,7 | 6,01 |
| | A0A0W4T739 | Conjugal transfer protein OS=Salmonella enterica OX=28901 GN=IN95_04500 PE=4 SV=1 - [A0A0W4T739_SALER] - R0127 - **TrhH** | 59,46 | 39,92 | 4 | 13 | 13 | 18 | 471 | 50,4 | 5,69 |
| | A0A3Z4T556 | Conjugal transfer protein TraN (Fragment) OS=Salmonella enterica I OX=59201 GN=DPC30_23900 PE=4 SV=1 - [A0A3Z4T556_SALET] - **TraN** | 4,87 | 3,34 | 6 | 2 | 2 | 2 | 927 | 102,1 | 5,63 |
| Δ-RSP (negative control) | A0A124DWT2 | Malate dehydrogenase OS=Salmonella enterica OX=28901 GN=mdh_2 PE=3 SV=1 - [A0A124DWT2_SALER] - **Mdh_2** | 12,6 | 11,25 | 70 | 2 | 2 | 3 | 311 | 32,2 | 6,11 |
| | G8I0D7 | Chloramphenicol acetyltransferase (Fragment) OS=Salmonella enterica subsp. enterica serovar Weltevreden OX=57743 GN=catA1 PE=4 SV=1 - [G8I0D7_SALET] - **CatA1** | 11,4 | 18,42 | 7 | 2 | 2 | 3 | 152 | 18 | 6,42 |
| | G5SCI6 | Uncharacterized protein (Fragment) OS=Salmonella enterica subsp. enterica serovar Wandsworth str. A4-580 OX=913086 GN=LTSEWAN_2946 PE=4 SV=1 - [G5SCI6_SALET] - **LTSEWAN_2946** | 11,03 | 72,22 | 16 | 2 | 2 | 3 | 36 | 3,9 | 4,61 |
| | A0A0V2F2B3 | 30S ribosomal protein S16 OS=Salmonella enterica subsp. enterica serovar Newport str. S09097 OX=1243581 GN=rpsP PE=3 SV=1 - [A0A0V2F2B3_SALNE] - **LFZ31_130775** | 9,47 | 32,93 | 10 | 2 | 2 | 3 | 82 | 9,2 | 10,55 |
| | A0A447MAR3 | Osmotically inducible protein OsmY OS=Salmonella bongori OX=54736 GN=osmY_1 PE=4 SV=1 - [A0A447MAR3_SALBN] - **OsmY_1** | 6,64 | 24,74 | 51 | 2 | 2 | 2 | 97 | 10,2 | 7,25 |
| | G5Q3W0 | Isocitrate dehydrogenase OS=Salmonella enterica subsp. enterica serovar Montevideo str. S5-403 OX=913242 GN=LTSEMON_2708 PE=4 SV=1 - [G5Q3W0_SALMO] - **LTSEMON_2708** | 6,57 | 15,17 | 87 | 3 | 3 | 3 | 290 | 31,7 | 5,24 |
| | A0A241SC82 | Anti-sigma-28 factor FlgM OS=Salmonella enterica subsp. salamae serovar 55:k:z39 str. 1315K OX=1243602 GN=LFZ47_10305 PE=4 SV=1 - [A0A241SC82_SALER] - **FlgM** | 6,51 | 22,68 | 20 | 2 | 2 | 3 | 97 | 10,6 | 9,41 |
| | A0A447U4Q8 | Zinc/cadmium-binding protein OS=Salmonella enterica I OX=59201 GN=zinT PE=4 SV=1 - [A0A447U4Q8_SALET] - **ZinT** | 6,39 | 13,95 | 88 | 2 | 2 | 2 | 172 | 19,9 | 8,59 |
| | A0A3R0EZK2 | UPF0265 protein YeeX OS=Salmonella enterica I OX=59201 GN=yeeX PE=3 SV=1 - [A0A3R0EZK2_SALET] - **YeeX** | 5,99 | 38,32 | 11 | 2 | 2 | 2 | 107 | 12,6 | 9,11 |
| | G5Q5T5 | Cysteine synthase (Fragment) OS=Salmonella enterica subsp. enterica serovar Montevideo str. S5-403 OX=913242 GN=LTSEMON_3543 PE=4 SV=1 - [G5Q5T5_SALMO] - **LTSEMON_3543** | 5,63 | 8,75 | 57 | 2 | 2 | 2 | 240 | 25,6 | 5,12 |
| | A0A379T767 | Enolase OS=Salmonella enterica subsp. arizonae OX=59203 GN=eno_3 PE=4 SV=1 - [A0A379T767_SALER] - **Eno_3** | 5,44 | 17,7 | 36 | 3 | 3 | 3 | 226 | 24,2 | 4,98 |
| | A0A379T547 | 50S ribosomal subunit protein L2 OS=Salmonella enterica subsp. arizonae OX=59203 GN=rplB_2 PE=4 SV=1 - [A0A379T547_SALER] - **RplB_2** | 5,43 | 19,9 | 18 | 3 | 3 | 4 | 196 | 21,4 | 10,87 |
| | A0A3S4FXF4 | Major type 1 subunit fimbrin (Pilin) OS=Salmonella enterica I OX=59201 GN=fimA_3 PE=4 SV=1 - [A0A3S4FXF4_SALET] - **FimA** | 5,24 | 14,56 | 123 | 2 | 2 | 2 | 158 | 16,2 | 4,97 |
| | G5LQ45 | Flagellar hook-associated protein 1 OS=Salmonella enterica subsp. enterica serovar Alachua str. R6-377 OX=913241 GN=flgK PE=3 SV=1 - [G5LQ45_SALET] - **FlgK** | 4,63 | 4,04 | 61 | 2 | 2 | 2 | 545 | 58,2 | 4,97 |
| | A0A3S5YH88 | 30S ribosomal protein S20 OS=Salmonella enterica subsp. arizonae serovar 18:z4,z23:- str. CVM N26626 OX=1395119 GN=P298_01775 PE=4 SV=1 - [A0A3S5YH88_SALER] - **P298_01775** | 4,58 | 31,15 | 10 | 2 | 2 | 2 | 61 | 6,7 | 10,36 |
| | A0A447NX75 | 5-methyltetrahydropteroyltriglutamate--homocysteine methyltransferase OS=Salmonella enterica subsp. salamae OX=59202 GN=metE_2 PE=3 SV=1 - [A0A447NX75_SALER] - **MetE_2** | 3,58 | 6,49 | 202 | 2 | 2 | 2 | 462 | 52 | 7,75 |
| | A0A3S5DC79 | Branched-chain amino acid aminotransferase OS=Salmonella enterica subsp. enterica serovar Sanjuan OX=1160765 GN=ilvE_1 PE=4 SV=1 - [A0A3S5DC79_SALET] - **IlvE_1** | 2,98 | 17,31 | 59 | 2 | 2 | 2 | 104 | 11,7 | 5,91 |

**Table S1**. Proteins copurifying with the RSP protein in an immunoprecipitation experiment. The blue label corresponds to strain SL1344 (R27 RSP-Flag). The yellow label corresponds to strain SL1344 (R27 Δ*rsp*) (negative control).

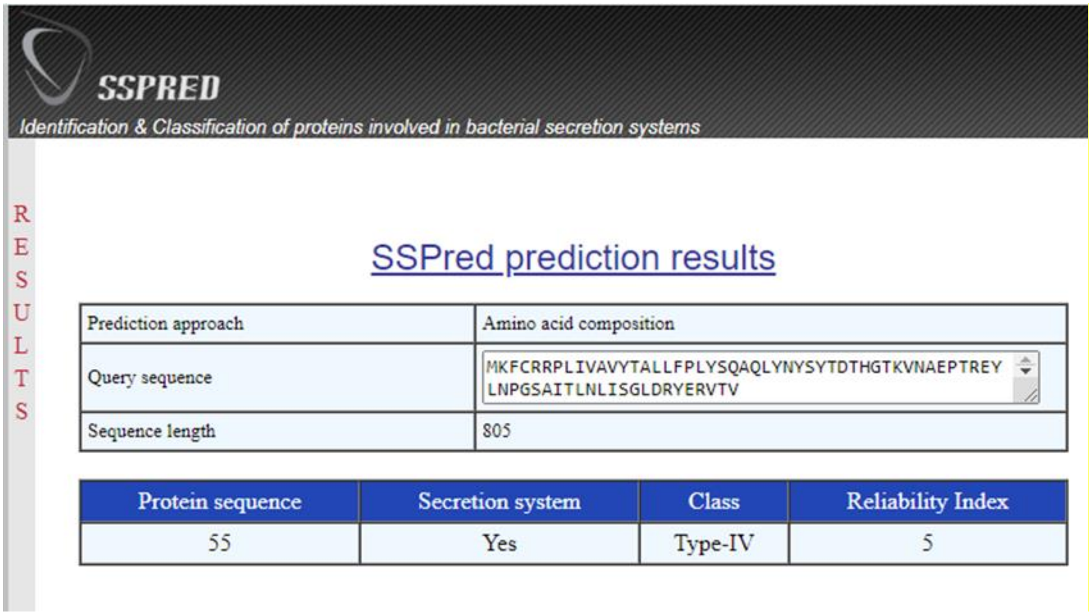


**Figure S1.** SSPRED algorithm prediction for RSP2 protein. Prediction approach was based on the RSP2 protein amino acid composition.

| Description | Incompatibility group | Accession number (NCBI) |
| --- | --- | --- |
| Escherichia coli isolate MSB1_9D-sc-2280338 plasmid 2 | IncHI1 | NZ_LR890350.1 |
| Escherichia coli O18ac:H14 strain 873.10 plasmid unnamed1 | IncHI1 | NZ_CP061755.12 |
| Escherichia coli strain ARL09/232 plasmid pCO_Eco4457-1 | IncHI1 | NZ_CP049968.13 |
| Escherichia coli strain ST410 plasmid p2189-NDM | IncHI1 | NZ_CP029631.14 |
| Salmonella enterica subsp. enterica serovar Typhi plasmid R27 | IncHI1 | NC_002305.15 |
| Salmonella enterica subsp. enterica serovar Derby strain CVM 30155 plasmid p30155-1 | IncHI1 | NZ_CP053049.16 |
| Escherichia coli strain RW7-1 plasmid pRW7-1_235k_tetX | IncFIA (HI1), IncHI1A, IncHI1B (R27), IncX1 | NZ_MT219825.17 |
| Escherichia coli strain RT18-1 plasmid pRT18-1_294k_tetX | IncFIB (K), IncFIA (HI1), IncHI1A, IncHI1B (R27) | NZ_MT219824.18 |
| Escherichia coli strain SY3626_hybrid plasmid pSY3626_190k_tetX | IncFIB (K), IncFIA (HI1), IncHI1A, IncHI1B (R27) | NZ_CP059284.19 |
| Escherichia coli strain SY3626C1 plasmid pSY3626C1_315k | IncFIB (K), IncFIA (HI1), IncHI1A, IncHI1B (R27) | NZ_CP059044.110 |
| Escherichia coli strain SY3626C5 plasmid pSY3626C1_229k | IncFIA (HI1), IncHI1A, IncHI1B (R27) | NZ_CP058949.111 |
| Escherichia coli strain SY3626 plasmid pSY3626_190k_tetX | IncFIA (HI1), IncHI1A, IncHI1B (R27) | NZ_JABXOE010000002.112 |
| Escherichia coli strain 2019XSD11-TC2 plasmid p2019XSD11-TC2-284 | IncFIA, IncFII, IncHI1A, IncHI1B | NZ_MN101858.113 |
| Escherichia coli strain 2019XSD11 plasmid p2019XSD11-190 | IncFIA, IncHI1A, IncHI1B | NZ_MN101856.114 |
| Escherichia coli strain 14OD0056 plasmid p14ODMR | IncHI1, IncFIA | NZ_MG904992.115 |
| Escherichia coli strain T28R plasmid pT28R-1 | IncFIA (HI1), IncHI1A, IncHI1B (R27) | NZ_CP049354.116 |
| Escherichia coli strain T16R plasmid pT16R-1 | IncFIA (HI1), IncHI1A, IncHI1B (R27) | NZ_CP046717.117 |
| Escherichia coli strain 1919D62 plasmid p1919D62-1 | IncFIA (HI1), IncHI1A, IncHI1B (R27) | NZ_CP046007.118 |
| Escherichia coli strain 1919D3 plasmid p1919D3-1 | IncFIA (HI1), IncHI1A, IncHI1B (R27) | NZ_CP046004.119 |
| Escherichia coli strain YPE10 plasmid pYPE10-190k-tetX4 | IncFIA (HI1), IncHI1A, IncHI1B (R27) | NZ_CP041449.120 |
| Escherichia coli strain YSP8-1 plasmid pYSP8-1 | IncHI1 | NZ_CP037911.121 |
| Escherichia coli strain CP53 plasmid pCP53-mcr | IncFIA, IncHI1A, IncHI1B, IncN | NZ_CP033094.122 |
| Salmonella enterica subsp. enterica serovar Typhi str. CT18 plasmid pHCM1 | IncHI1 | NC_003384.124 |
| Salmonella enterica strain GX1006 plasmid pSal21GXH-tetX4 | IncFIA (HI1), IncHI1A, IncHI1B (R27) | NZ_CP060586.125 |
| Salmonella enterica subsp. enterica serovar Typhimurium strain R18.1078 plasmid pR18.1078_p247k | IncFIA (HI1), IncHI1A, IncHI1B (R27) | NZ_CP065568.126 |
| Salmonella enterica subsp. enterica serovar Typhi strain 311189_206186 plasmid pHCM1 | IncHI1 | NZ_CP029939.128 |
| Salmonella enterica subsp. enterica serovar Enteritidis strain 81-1705 plasmid pSE81-1705-1 | IncFIA (HI1), IncHI1A, IncHI1B (R27) | NZ_CP018652.148 |
| Salmonella enterica subsp. enterica serovar Enteritidis strain 81-1706 plasmid pSE81-1706 | IncFIA (HI1), IncFIB (S), IncFII (S), IncHI1A, IncHI1B (R27) | NZ_CP018656.151 |

| | | |
|---|---|---|
| Escherichia coli strain SD134209 plasmid pSD134209-1 | IncFIA (HI1), IncHI1A, IncHI1B (R27) | NZ_CP029690.152 |
| Salmonella enterica subsp. enterica serovar Saintpaul strain 4695 plasmid p4695_blaTEM-1B | IncFIA (HI1), IncHI1A, IncHI1B (R27) | NZ_CP062979.253 |
| Salmonella enterica subsp. enterica serovar Typhimurium strain 21G7 isolate B71 plasmid pB71 | IncHI1 | NZ_KP899806.154 |
| Escherichia coli strain 4M9F plasmid p4M9F | IncFIA (HI1)/IncHI1A/IncHI1B (R27) | NZ_MN256759.155 |
| Escherichia coli strain 4M8F plasmid p4M8F | IncFIA (HI1), IncFIB, IncHI1A, IncHI1B (R27),IncY | NZ_MN256758.156 |
| Escherichia coli strain 4M18F plasmid p4M18F | IncFIA (HI1), IncFIB, IncHI1A, IncHI1B (R27),IncY | NZ_MN256757.157 |
| Klebsiella pneumoniae strain KP14812 plasmid pKP14812-MCR-1 | IncHI1 | NZ_MH733010.158 |
| Escherichia coli strain CP131_Sichuan plasmid pCP131-IncHI1 | IncHI1 | NZ_CP053721.159 |
| Escherichia coli strain EC2 plasmid pEC2-4 | IncHI1 | NZ_CP016184.160 |
| Salmonella enterica strain SA20030575 plasmid pSA20030575.1 | IncHI1A, IncHI1B (R27) | NZ_CP030182.161 |
| Salmonella enterica subsp. enterica serovar Saintpaul strain SGB23 plasmid pSGB23 | IncHI1 | NZ_CP023167.162 |
| Salmonella enterica subsp. enterica serovar Typhimurium strain CVM 28321-a plasmid p28321a-1 | IncFIA-IncHI1A-InHI1B-IncQ | NZ_CP053053.163 |
| Salmonella enterica subsp. enterica serovar Typhimurium strain CVM 24362 plasmid p24362-1 | IncFIA (HI1), IncHI1A, IncHI1B (R27), IncQ1 | NZ_CP051379.164 |
| Escherichia coli strain 100063-3 plasmid p100063-3 | IncFIA (HI1), IncHI1A, IncHI1B (R27), IncQ1 | NZ_MT586609.165 |
| Escherichia coli strain 99063 plasmid p99063 | IncFIA (HI1), IncHI1A, IncHI1B (R27), IncQ1 | NZ_MT586606.167 |
| Escherichia coli strain 97974-2T plasmid p97974-2T | IncFIA (HI1), IncHI1A, IncHI1B (R27), IncQ1 | NZ_MT586605.168 |
| Escherichia coli strain 10068 plasmid p10068 | IncFIA (HI1), IncHI1A, IncHI1B (R27), IncQ1 | NZ_MT586604.169 |
| Escherichia coli strain 15S04829-4 plasmid p15S04829-4 | IncFIA (HI1), IncHI1A, IncHI1B (R27), IncQ1 | NZ_MT586603.170 |
| Escherichia coli strain 15S04714-1 plasmid p15S04714-1 | IncFIA (HI1), IncHI1A, IncHI1B (R27), IncQ1 | NZ_MT586601.171 |
| Escherichia coli strain 99783-3T plasmid p99783-3T | IncFIA (HI1), IncHI1A, IncHI1B (R27), IncQ1 | NZ_MT586607.172 |
| Escherichia coli strain 15S04779-4 plasmid p15S04779-4 | IncFIA (HI1), IncHI1A, IncHI1B (R27), IncQ1 | NZ_MT586602.173 |
| Salmonella enterica subsp. enterica serovar Typhimurium strain 21G5 isolate 109-9 plasmid p109/9 | IncHI1 | NZ_KP899805.174 |

| | | |
|---|---|---|
| Salmonella enterica subsp. enterica serovar Typhimurium strain 21G6 isolate F8475 plasmid pF8475 | IncHI1 | NZ_KP899804.175 |
| Klebsiella michiganensis strain RHB20-C01 plasmid pRHB20-C01_2 | IncFIA (HI1), IncHI1A, IncHI1B (R27), IncQ1 | NZ_JABXTO010000002.176 |
| Escherichia coli plasmid pGD27-31 | IncFIA (HI1), IncHI1A, IncHI1B (R27) | NZ_MN232190.177 |
| Klebsiella michiganensis strain RHB20-C02 plasmid pRHB20-C02_2 | IncFIA (HI1), IncHI1A, IncHI1B (R27), IncQ1 | NZ_CP058213.178 |
| Salmonella enterica strain SRC27 plasmid pSRC27-H | IncHI1 | NZ_CP058810.179 |
| Escherichia coli plasmid pEQ2 | IncHI1 | NC_023277.280 |
| Escherichia coli plasmid pEQ1 | IncHI1 | NC_023289.281 |
| Escherichia coli strain 3498 plasmid p3498 | IncFIA (HI1), IncHI1A, IncHI1B (R27), IncQ1 | NZ_MG948335.182 |
| Escherichia coli strain H226B plasmid pH226B | IncHI1 | NZ_KX129784.183 |
| Escherichia coli strain 1454 plasmid RCS78_p | IncFIA (HI1), IncHI1A, IncHI1B (R27), IncQ1 | NZ_LT985296.184 |
| Salmonella enterica subsp. enterica serovar strain PNCS000211 plasmid p10-3184.1 | IncFIA (HI1), IncHI1A, IncHI1B (R27) | NZ_CP039717.188 |
| Salmonella enterica subsp. enterica serovar strain PNCS014846 plasmid p08-4425.1 | IncFIA (HI1), IncHI1A, IncHI1B (R27) | NZ_CP039559.189 |
| Escherichia coli O111:H- str. 11128 plasmid pO111_1 | IncHI1 | NC_013365.190 |
| Salmonella enterica subsp. enterica serovar Choleraesuis plasmid pMAK1 | IncHI1 | NC_009981.191 |
| Escherichia coli strain H9Ecoli plasmid p1-H9 | p0111 | NZ_CP029181.193 |
| Escherichia coli strain ECCHD184 plasmid pTB211 | p0111, IncFIB (AP001918) | NZ_CP033251.194 |
| Escherichia coli plasmid p16EC-p0111 | p0111, IncN-type | NZ_MN086777.195 |
| Escherichia coli strain ECCRA-119 plasmid pTB202 | p0111, IncN-type | NZ_CP029244.196 |
| Escherichia coli strain EC2_1 plasmid pEC2_1-4 | IncHI, IncF | NZ_CP016183.197 |

**Table S2.** Distribution of the RSP2 protein among bacterial plasmids.

```
pKAZ3               ----MINFKPKIPAMLGALAVLTAGAAHAELLEYTFKAPDGTQRSLTPNANYANPTGNIS 56
pMS6198A            ----MINFKPKIPAMLGALAVLTAGAAHAELLEYTFKAPDGTQRSLTPNANYANPTGNIS 56
pKp55               MENAMINFKPKIPAMLGALAVLTAGAAHAELLEYTFKAPDGTQRSLTPNANYANPTGNIS 60
pEc19               MENAMINFKPKIPAMLGALAVLTAGAAHAELLEYTFKAPDGTQRSLTPNANYANPTGNIS 60
pAPEC1990           ----MINFKPKIPAMLGALAVLTAGAAHAELLEYTFKAPDGTQRSLTPNANYANPTGNIS 56
pNDM-KN             ----MINFKPKIPAMLGALAVLTAGAAHAELLEYTFKAPDGTQRSLTPNANYANPTGNIS 56
pUMNK88             ----MINFKPKIPAMLGALAVLTAGAAHAELLEYTFKAPDGTQRSLTPNANYANPTGNIS 56
pCf587              ----MINFKPKLPALLGALAVLTAGSAHAELLEYTFKAPDGAQRSLPPNANYANPTGNVS 56
                        *******:**:**********:***************:**** ***********:*

pKAZ3               FALSAGIDRKVKISVIRSDGTVVSTATSHLLGATDRITVGGKSYYGAELQLPAPSEGVYK 116
pMS6198A            FALSAGIDRKVKISVLRSDGTVVSTATSHLLGATDRITVGGKSYYGAELQLPAPVGGAYT 116
pKp55               FALSAGIDRKVKISVLRSDGTVVSTATSHLLGATDRITVGGKSYYGAELQLPAPVGGAYT 120
pEc19               FALSAGIDRKVKISVLRSDGTVVSTATSHLLGATDRITVGGKSYYGAELQLPAPVGGAYT 120
pAPEC1990           FALSAGIDRKVKISVLRSDGTVVSTATSHLLGATDRITVGGKSYYGAELQLPAPVGGAYT 116
pNDM-KN             FALSAGIDRKVKISVLRSDGTVVSTATSHLLGATDRITVGGKSYYGAELQLPAPVGGAYT 116
pUMNK88             FALSAGIDRKVKISVLRSDGTVVSTATSHLLGATDRITVGGKSYYGAELQLPAPVGGAYT 116
pCf587              FALSAGIDRKVKVSVRRSDGTVVSTATSHLLGATDRITVGGKSYYGAELQLAAPAAGSYT 116
                    ************:** *********************************** **  * *.

pKAZ3               LRAEILASDGSSVQTDEYPLQVDTTAPSLANVTV----------KGEWNRALSDGTLLRG 166
pMS6198A            IRAEILASDGSTVQTDEYPLTVDVTPPTYSSLAPVYSNYGQVTSGDVWKLGLGGSE--DN 174
pKp55               IRAEILASDGSTVQTDEYPLTVDVTPPTYSSLAPVYSNYGQVTSGDVWKLGLGGSE--DN 178
pEc19               IRAEILASDGSTVQTDEYPLTVDVTPPTYSSLAPVYSNYGQVTSGDVWKLGLGGSE--DN 178
pAPEC1990           IRAEILASDGSTVQTDEYPLTVDVTPPTYSSLAPVYSNYGQVTSGDVWKLGLGGSE--DN 174
pNDM-KN             IRAEILASDGSTVQTDEYPLTVDVTPPTYSSLAPVYSNYGQVTSGDVWKLGLGGSE--DN 174
pUMNK88             IRAEILASDGSTVQTDEYPLTVDVTPPTYSSLAPVYSNYGQVTSGDVWKLGLGGSE--DN 174
pCf587              IRAEILASDGSTVQSDDYPLTVDVTPPKYTSLAPVYGTYGQVISGDVWKLGTGGAE--EN 174
                    :**********:**:*:*** **.* *. :.::            . *: . ...    .

pKAZ3               PNRFSGIDVSASDAGSSVSSIQAYAVDSKGKRSPVVSVNYANGQGQL--LNWSTVFPN-- 222
pMS6198A            AFLLSGIS-----DESPIKGVKAKLYRQDGSLYKDVSVNYDDANGQARQSFQSGFFPASD 229
pKp55               AFLLSGIS-----DESPIKGVKAKLYRQDGSLYKDVSVNYDDANGQARQSFQSGFFPASD 233
pEc19               AFLLSGIS-----DESPIKGVKAKLYRQDGSLYKDVSVNYDDANGQARQSFQSGFFPASD 233
pAPEC1990           AFLLSGIS-----DESPIKGVKAKLYRQDGSLYKDVSVNYDDANGQARQSFQSGFFPASD 229
pNDM-KN             AFLLSGIS-----DESPIKGVKAKLYRQDGSLYKDVSVNYDDANGQARQSFQSGFFPASD 229
pUMNK88             AFLLSGIS-----DESPIKGVKAKLYRQDGSLYKDVSVNYDDANGQARQSFQSGFFPASD 229
pCf587              AFLISGIS-----DDSPVKEVKAKLYRQDGSLYKDVSVNYDDANKQARQYFESGFFPASD 229
                       :***.       * :. ::*    ..*.    ***** :.: *      * .**

pKAZ3               GEDLYTLHFEALDKAGNKGSIA-YPIAWDSVGAKSGENPEPVAVYDPKNPQASTFKVNGQ 281
pMS6198A            LDEVFTLQFEISDSAGNSYLSPRQKVMFDSI---TNAPSAPFGVYDPSST-----NNLGP 281
pKp55               LDEVFTLQFEISDSAGNSYLSPRQKVMFDSI---TNAPSAPFGVYDPSST-----NNLGP 285
pEc19               LDEVFTLQFEISDSAGNSYLSPRQKVMFDSI---TNAPSAPFGVYDPSST-----NNLGP 285
pAPEC1990           LDEVFTLQFEISDSAGNSYLSPRQKVMFDSI---TNAPSAPFGVYDPSST-----NNLGP 281
pNDM-KN             LDEVFTLQFEISDSAGNSYLSPRQKVMFDSI---TNAPSAPFGVYDPSST-----NNLGP 281
pUMNK88             LDEVFTLQFEISDSAGNSYLSPRQKVMFDSI---TNAPSAPFGVYDPSST-----NNLGP 281
pCf587              LDEVFTLQFQLSDSAGNSYLSPPQKVMFDNL---TNAPSSPFGVYDPASS-----STLGP 281
                     ::::**:*:  *.***.       : :*.:   :.    *..**** .      .  *

pKAZ3               ALSGFAPYQNGMTIYSDTYRVLYRIPKTNSYPSSPYGAQSGNWCDYKNCI--------EG 333
pMS6198A            GLTGFVAYTEGMTVKTNPIKLAWRVPRDN-------------WHEYREGGINMTNALGEM 328
pKp55               GLTGFVAYTEGMTVKTNPIKLAWRVPRDN-------------WHEYREGGINMTNALGEM 332
pEc19               GLTGFVAYTEGMTVKTNPIKLAWRVPRDN-------------WHEYREGGINMTNALGEM 332
pAPEC1990           GLTGFVAYTEGMTVKTNPIKLAWRVPRDN-------------WHEYREGGINMTNALGEM 328
pNDM-KN             GLTGFVAYTEGMTVKTNPIKLAWRVPRDN-------------WHEYREGGINMTNALGEM 328
pUMNK88             GLTGFVAYTEGMTVKTNPIKLAWRVPRDN-------------WHEYREGGINMTNALGEM 328
pCf587              GLSGFVAYKAGMTVKTNPIKLAWRVPRNN-------------WHEYREGGINMVNSLGEM 328
                    .*:**. *  ***: ::  :: :*:*: *             * :*::          *

pKAZ3               NIIAEDGTYDYRQTQADVFQQHGTKTKSFIIYDWVMNS-L--GNTSVSVKADPSVSLAPI 390
pMS6198A            SKVGEDASYVYLVTTAPYGNTDGN------YWRWVNFGQWGGGGIAYNLTLSPSAPKSPR 382
pKp55               SKVGEDASYVYLVTTAPYGNTDGN------YWRWVNFGQWGGGGIAYNLTLSPSAPKSPR 386
pEc19               SKVGEDASYVYLVTTAPYGNTDGN------YWRWVNFGQWGGGGIAYNLTLSPSAPKSPR 386
pAPEC1990           SKVGEDASYVYLVTTAPYGNTDGN------YWRWVNFGQWGGGGIAYNLTLSPSAPKSPR 382
pNDM-KN             SKVGEDASYVYLVTTAPYGNTDGN------YWRWVNFGQWGGGGIAYNLTLSPSAPKSPR 382
pUMNK88             SKVGEDASYVYLVTTAPYGNTDGN------YWRWVNFGQWGGGGIAYNLTLSPSAPKSPR 382
pCf587              TKVGEDGNYVYLVTTAPYGNTDGN------YWRWVNFGQWGGGVIAYDLSLSPSAPQSPK 382
                    . :.**..* *  * *   : .*.       : **  .    *  : .:. .**.  :*

pKAZ3               GKYVEYLRDDG-------QWVRGETINMSSVNHYKKLRFHVEPRPYAQELWGSWLPTTKI 443
pMS6198A            LLGVDYNYSDIGWSSFYRYWVNNSVLP----VTVSSIRVKVEPRPYVQTAVHRG--SC-- 434
pKp55               LLGVDYNYSDIGWSSFYRYWVNNSVLP----VTVSSIRVKVEPRPYVQTAVHRG--SC-- 438
pEc19               LLGVDYNYSDIGWSSFYRYWVNNSVLP----VTVSSIRVKVEPRPYVQTAVHRG--SC-- 438
pAPEC1990           LLGVDYNYSDIGWSSFYRYWVNNSVLP----VTVSSIRVKVEPRPYVQTAVHRG--SC-- 434
pNDM-KN             LLGVDYNYSDIGWSSFYRYWVNNSVLP----VTVSSIRVKVEPRPYVQTAVHRG--SC-- 434
pUMNK88             LLGVDYNYSDIGWSSFYRYWVNNSVLP----VTVSSIRVKVEPRPYVQTAVHRG--SC-- 434
```

```
pCf587         LLGVDYNYSDIGWSSMYRYWVDSSVLP----VTVSSIRVKVEPRPYVQTAVHRG--SC-- 434
                  *:*  .*         ** ...:        ..:*.:******.*        :

pKAZ3          PANASYAEVDTDISFSGRSCSWPGYWSRPEGKPELPSDR--------------IGAT--F 487
pMS6198A       -----------EIPVGQDSCVIANSFTMAKGTTGYVHDNATVFNPDKSLRSNPLWAEVNW 483
pKp55          -----------EIPVGQDSCVIANSFTMAKGTTGYVHDNATVFNPDKSLRSNPLWAEVNW 487
pEc19          -----------EIPVGQDSCVIANSFTMAKGTTGYVHDNATVFNPDKSLRSNPLWAEVNW 487
pAPEC1990      -----------EIPVGQDSCVIANSFTMAKGTTGYVHDNATVFNPDKSLRSNPLWAEVNW 483
pNDM-KN        -----------EIPVGQDSCVIANSFTMAKGTTGYVHDNATVFNPDKSLRSNPLWAEVNW 483
pUMNK88        -----------EIPVGQDSCVIANSFTMAKGTTGYVHDNATVFNPDKSLRSNPLWAEVNW 483
pCf587         -----------EVPVGQDSCVIANSFTMSKGTTGYIHDNATVFNADRSLSSNPLWAEVNW 483
                          :: ..  **   . ::  :*.     *.              : *   :

pKAZ3          CYDLNPPEIAGLERNGRIFKAHFREPDS---FDGWGV-NQWVISNNSSASAITSTGEERP 543
pMS6198A       NDQHYPQLSQQFDQNSKVFTLFVNQPGRGAYFDRLRLRSAWIEDSKG--NKLS------- 534
pKp55          NDQHYPQLSQQFDQNSKVFTLFVNQPGRGAYFDRLRLRSAWIEDSKG--NKLS------- 538
pEc19          NDQHYPQLSQQFDQNSKVFTLFVNQPGRGAYFDRLRLRSAWIEDSKG--NKLS------- 538
pAPEC1990      NDQHYPQLSQQFDQNSKVFTLFVNQPGRGAYFDRLRLRSAWIEDSKG--NKLS------- 534
pNDM-KN        NDQHYPQLSQQFDQNSKVFTLFVNQPGRGAYFDRLRLRSAWIEDSKG--NKLS------- 534
pUMNK88        NDQHYPQLSQQFDQNSKVFTLFVNQPGRGAYFDRLRLRSAWIEDSKG--NKLS------- 534
pCf587         NDQHYPQLSQQYDQSNKVFTLFVNQPGRGAYFDRLRLRSAWIEDSKG--NKLS------- 534
                 :  *      ::..::*. ...:*.    **   : . *: ..:.  . ::

pKAZ3          LSRTEVLRSSTNDWFFTFSAENLSEGTYTGISLVAKDAFGNEVKQVFTGSQYAMSIDNSA 603
pMS6198A       -PTGGLIANNWENYTYQWDLKTLPEGQYSLVAAA-EEMHG----PLTRQPMFQITSDRTP 588
pKp55          -PTGGLIANNWENYTYQWDLKTLPEGQYSLVAAA-EEMHG----PLTRQPMFQITSDRTP 592
pEc19          -PTGGLIANNWENYTYQWDLKTLPEGQYSLVAAA-EEMHG----PLTRQPMFQITSDRTP 592
pAPEC1990      -PTGGLIANNWENYTYQWDLKTLPEGQYSLVAAA-EEMHG----PLTRQPMFQITSDRTP 588
pNDM-KN        -PTGGLIANNWENYTYQWDLKTLPEGQYSLVAAA-EEMHG----PLTRQPMFQITSDRTP 588
pUMNK88        -PTGGLIANNWENYTYQWDLKTLPEGQYSLVAAA-EEMHG----PLTRQPMFQITSDRTP 588
pCf587         -PTGGLIANNWENYTYQWDLKTLPEGQYSLVAAA-EEMHG----PLTRQPMFQITSDKTA 588
                    :: .. ::: : :. :.* ** *: :: . :: .*     :     : :: *.:

pKAZ3          PTLTVSISDGAPIQSLDDVVITLTDTADPSPKLTSIALVGGPADDKVQLSWREESKGRFR 663
pMS6198A       PTMTLSVADGAAIQTLDDVVITLADAIDPSPKLTSIALVGGPANDKVQLSWREESKGRFR 648
pKp55          PTMTLSVADGAAIQTLDDVVITLADAIDPSPKLTSIALVGGPANDKVQLSWREESKGRFR 652
pEc19          PTMTLSVADGAAIQTLDDVVITLADAIDPSPKLTSIALVGGPANDKVQLSWREESKGRFR 652
pAPEC1990      PTMTLSVADGAAIQTLDDVVITLADAIDPSPKLTSIALVGGPANDKVQLSWREESKGRFR 648
pNDM-KN        PTMTLSVADGAAIQTLDDVVITLADAIDPSPKLTSIALVGGPANDKVQLSWREESKGRFR 648
pUMNK88        PTMTLSVADGAAIQTLDDVVITLADAIDPSPKLTSIALVGGPANDKVQLSWREESKGRFR 648
pCf587         PTLTISVADGAAIQTLDDVVITLADAIDPSPKLTSIALVGGPANDKVQLSWREESKGRFR 648
               **:*:*::*** **:********:*: ****************:****************

pKAZ3          LEYPVMFPSLKEGESYTLTVSGEDAQGNAVQKAVGFEYKPRQVMLADGMDGKVMVPAVTH 723
pMS6198A       LEYPVMFPSLKEGESYTLTVSGEDAQGNAVQKAVGFEYKPRQVMLADGMDGKVMVPAVTH 708
pKp55          LEYPVMFPSLKEGESYTLTVSGEDAQGNAVQKAVGFEYKPRQVMLADGMDGKVMVPAVTH 712
pEc19          LEYPVMFPSLKEGESYTLTVSGEDAQGNAVQKAVGFEYKPRQVMLADGMDGKVMVPAVTH 712
pAPEC1990      LEYPVMFPSLKEGESYTLTVSGEDAQGNAVQKAVGFEYKPRQVMLADGMDGKVMVPAVTH 708
pNDM-KN        LEYPVMFPSLKEGESYTLTVSGEDAQGNAVQKAVGFEYKPRQVMLADGMDGKVMVPAVTH 708
pUMNK88        LEYPVMFPSLKEGESYTLTVSGEDAQGNAVQKAVGFEYKPRQVMLADGMDGKVMVPAVTH 708
pCf587         LEYPVMFPSLKEGESYRLTVSGADAQGNAVKKVVGFEYKPRQVMLADGMDGKVMVPAVTH 708
               **************** ***** *******:*.***************************

pKAZ3          EFVHADGKRIIETKPLTLSDGAVVTGSYDVFATLRSDAKVPLVVNGVRIEPGQTMGIMSQ 783
pMS6198A       EFVHADGKRIIETKPLTLSDGAVVTGSYDVFATLRSDAKVPLVVNGVRIEPGQTMGIMSQ 768
pKp55          EFVHADGKRIIETKPLTLSDGAVVTGSYDVFATLRSDAKVPLVVNGVRIEPGQTMGIMSQ 772
pEc19          EFVHADGKRIIETKPLTLSDGAVVTGSYDVFATLRSDAKVPLVVNGVRIEPGQTMGIMSQ 772
pAPEC1990      EFVHADGKRIIETKPLTLSDGAVVTGSYDVFATLRSDAKVPLVVNGVRIEPGQTMGIMSQ 768
pNDM-KN        EFVHADGKRIIETKPLTLSDGAVVTGSYDVFATLRSDAKVPLVVNGVRIEPGQTMGIMSQ 768
pUMNK88        EFVHADGKRIIETKPLTLSDGAVVTGSYDVFATLRSDAKVPLVVNGVRIEPGQTMGIMSQ 768
pCf587         EFVHADGKRIIETKPLTLSDGAIVTGSYDVFATLRSDAKVPLVVNGVRIEPGQTMGIMSQ 768
               **********************:*************************************

pKAZ3          HDFGASGGRLSIPVKPAVPDVVGSSSLLVMTSAPNSPILVVDINTWKGTAKLSAESWTIR 843
pMS6198A       HDFGASGGRLSIPVKPAVPDVVGSSSLLVMTSAPNSPILVVDINTWKGTAKLSAESWTIR 828
pKp55          HDFGASGGRLSIPVKPAVPDVVGSSSLLVMTSAPNSPILVVDINTWKGTAKLSAESWTIR 832
pEc19          HDFGASGGRLSIPVKPAVPDVVGSSSLLVMTSAPNSPILVVDINTWKGTAKLSAESWTIR 832
pAPEC1990      HDFGASGGRLSIPVKPAVPDVVGSSSLLVMTSAPNSPILVVDINTWKGTAKLSAESWTIR 828
pNDM-KN        HDFGASGGRLSIPVKPAVPDVVGSSSLLVMTSAPNSPILVVDINTWKGTAKLSAESWTIR 828
pUMNK88        HDFGASGGRLSIPVKPAVPDVVGSSSLLVMTSAPNSPILVVDINTWKGTAKLSAESWTIR 828
pCf587         HDFGASGGRLSIPVKPAIPDVVGSSSLLVMTSAPNSPILVVDINTWKGAAKLSAESWTIR 828
               *****************:******************************:***********

pKAZ3          QVIDPVKIYALPESGVPCRFTTKEDVAMAADPIRDPVCLLQWDRTPDEAEQTTQDNNGMK 903
pMS6198A       QVIDPVKIYALPESGVPCRFTTKEDVAMAADPIRDPVCLLQWDRTPDEAEQTTQDNNGMK 888
pKp55          QVIDPVKIYALPESGVPCRFTTKEDVAMAADPIRDPVCLLQWDRTPDEAEQTTQDNNGMK 892
pEc19          QVIDPVKIYALPESGVPCRFTTKEDVAMAADPIRDPVCLLQWDRTPDEAEQTTQDNNGMK 892
```

```
pAPEC1990      QVIDPVKIYALPESGVPCRFTTKEDVAMAADPIRDPVCLLQWDRTPDEAEQTTQDNNGMK 888
pNDM-KN        QVIDPVKIYALPESGVPCRFTTKEDVAMAADPIRDPVCLLQWDRTPDEAEQTTQDNNGMK 888
pUMNK88        QVIDPVKIYALPESGVPCRFTTKEDVAMAADPIRDPVCLLQWDRTPDEAEQTTQDNNGMK 888
pCf587         QVIDPVKIYALPETGVPCRFTTKADVAMAADPIRDPVCLLQWDRTPDEAEQTTQDTNGMK 888
               *************:********* *******************************.****

pKAZ3          VAGLVGQAVSIGEQPVEYSLYLFSGDGSKVKVGSGSQNLTVTTAYGSVGYTPIDDIAQVN 963
pMS6198A       VAGLVGQAVSIGEQPVEYSLYLFSGDGSKVKVGSGSQNLTVTTAYGSVGYTPIDDIAQVN 948
pKp55          VAGLVGQAVSIGEQPVEYSLYLFSGDGSKVKVGSGSQNLTVTTAYGSVGYTPIDDIAQVN 952
pEc19          VAGLVGQAVSIGEQPVEYSLYLFSGDGSKVKVGSGSQNLTVTTAYGSVGYTPIDDIAQVN 952
pAPEC1990      VAGLVGQAVSIGEQPVEYSLYLFSGDGSKVKVGSGSQNLTVTTAYGSVGYTPIDDIAQVN 948
pNDM-KN        VAGLVGQAVSIGEQPVEYSLYLFSGDGSKVKVGSGSQNLTVTTAYGSVGYTPIDDIAQVN 948
pUMNK88        VAGLVGQAVSIGEQPVEYSLYLFSGDGSKVKVGSGSQNLTVTTAYGSVGYTPIDDIAQVN 948
pCf587         VAGLVGQAVSIGEQPVEYSLYLFSGDGSKVKVGSGSQNLTVTTAYGSVGYTPIDDIAQVN 948
               ************************************************************

pKAZ3          RVIEDFDVNFKQSKGPDCSITLSADRAKNEAANKAVGSASRTCLFEWQQIPDGLVQDPLS 1023
pMS6198A       RVIEDFDVNFKQSKGPDCSITLSADRAKKEAANKAVGSASRACLFEWQQIPDGLVQDPLS 1008
pKp55          RVIEDFDVNFKQSKGPDCSITLSADRAKKEAANKAVGSASRACLFEWQQIPDGLVQDPLS 1012
pEc19          RVIEDFDVNFKQSKGPDCSITLSADRAKKEAANKAVGSASRACLFEWQQIPDGLVQDPLS 1012
pAPEC1990      RVIEDFDVNFKQSKGPDCSITLSADRAKKEAANKAVGSASRACLFEWQQIPDGLVQDPLS 1008
pNDM-KN        RVIEDFDVNFKQSKGPDCSITLSADRAKKEAANKAVGSASRACLFEWQQIPDGLVQDPLS 1008
pUMNK88        RVIEDFDVNFKQSKGPDCSITLSADRAKKEAANKAVGSASRACLFEWQQIPDGLVQDPLS 1008
pCf587         RVIENFNVNLKQNKGPDCSITLSADRAKKEAASKGAGSASRTCLFEWQQIPDGLLQEQLS 1008
               ****:*:**:**.***************:***.*..*****:************:*: **

pKAZ3          ESPSLSGSLAENGDHPLGWRVSIFTRNGTRVTLNDETFNVEAVDPPAPTVELASDYNFKD 1083
pMS6198A       ESPSLSGSLASNGVHPLGWRVSIFTRNGTRVTLNDETFNVEAVDPPAPTVELASDYNFKD 1068
pKp55          ESPSLSGSLASNGVHPLGWRVSIFTRNGTRVTLNDETFNVEAVDPPAPTVELASDYNFKD 1072
pEc19          ESPSLSGSLASNGVHPLGWRVSIFTRNGTRVTLNDETFNVEAVDPPAPTVELASDYNFKD 1072
pAPEC1990      ESPSLSGSLASNGVHPLGWRVSIFTRNGTRVTLNDETFNVEAVDPPAPTVELASDYNFKD 1068
pNDM-KN        ESPSLSGSLASNGVHPLGWRVSIFTRNGTRVTLNDETFNVEAVDPPAPTVELASDYNFKD 1068
pUMNK88        ESPSLSGSLASNGVHPLGWRVSIFTRNGTRVTLNDETFNVEAVDPPAPTVELASDYNFKD 1068
pCf587         ESPSLSGSLASNGNHPLGWRVSIFTRNGTRVTLNDQTFNIEAVDPPAPTVELSSKFNFKD 1068
               **********.** *********************:***:************:*.:****

pKAZ3          NIYLVPMTGNYLGDAIINSERADLDIAISRNSDVLESETFTPGWGATNKVYRRINTDERA 1143
pMS6198A       NIYLVPMTGNYLGDAIINSERADLDIAISRNSDVLESETFTPGWGATNKVYRRINTDERA 1128
pKp55          NIYLVPMTGNYLGDAIINSERADLDIAISRNSDVLESETFTPGWGATNKVYRRINTDERA 1132
pEc19          NIYLVPMTGNYLGDAIINSERADLDIAISRNSDVLESETFTPGWGATNKVYRRINTDERA 1132
pAPEC1990      NIYLVPMTGNYLGDAIINSERADLDIAISRNSDVLESETFTPGWGATNKVYRRINTDERA 1128
pNDM-KN        NIYLVPMTGNYLGDAIINSERADLDIAISRNSDVLESETFTPGWGATNKVYRRINTDERA 1128
pUMNK88        NIYLVPMTGNYLGDAIINSERADLDIAISRNSDVLESETFTPGWGATNKVYRRINTDERA 1128
pCf587         NIYMVPMTGNYLGDAIISSERADLDIAISRNSGVLESETFAPGWGATNKVYRRINTDERA 1128
               ***:*************.**************.*******:*******************

pKAZ3          LWEETTYKVNAAYNKVPDVKTEVVYRAISAPSDSIRPIVEVKGDTAIDTQALPVRVLIRD 1203
pMS6198A       LWEETTYKVNAAYNKVPDVKTEVVYRAISAPSDSIRPIVEVKGDTAIDTQALPVRVLIRD 1188
pKp55          LWEETTYKVNAAYNKVPDVKTEVVYRAISAPSDSIRPIVEVKGDTAIDTQALPVRVLIRD 1192
pEc19          LWEETTYKVNAAYNKVPDVKTEVVYRAISAPSDSIRPIVEVKGDTAIDTQALPVRVLIRD 1192
pAPEC1990      LWEETTYKVNAAYNKVPDVKTEVVYRAISAPSDSIRPIVEVKGDTAIDTQALPVRVLIRD 1188
pNDM-KN        LWEETTYKVNAAYNKVPDVKTEVVYRAISAPSDSIRPIVEVKGDTAIDTQALPVRVLIRD 1188
pUMNK88        LWEETTYKVNAAYNKVPDVKTEVVYRAISAPSDSIRPIVEVKGDTAIDTQALPVRVLIRD 1188
pCf587         LWEETTYKVNAAYNKVPDVKTEAIYRAIAAPSDSIRPVVEVDGDNAIDTQALPVRVLIRD 1188
               **********************.:****:********:***.**.***************

pKAZ3          QYKPDGDYDANTMGVWKVRLIQQKAYNETVALTDYAEASNGEAQFSVDLSGVDTSSVRIA 1263
pMS6198A       QYKPDGDYDANTMGVWKVRLIQQKAYNETVALTDYAEASNGEAQFSVDLSGVDTSSVRIA 1248
pKp55          QYKPDGDYDANTMGVWKVRLIQQKAYNETVALTDYAEASNGEAQFSVDLSGVDTSSVRIA 1252
pEc19          QYKPDGDYDANTMGVWKVRLIQQKAYNETVALTDYAEASNGEAQFSVDLSGVDTSSVRIA 1252
pAPEC1990      QYKPDGDYDANTMGVWKVRLIQQKAYNETVALTDYAEASNGEAQFSVDLSGVDTSSVRIA 1248
pNDM-KN        QYKPDGDYDANTMGVWKVRLIQQKAYNETVALTDYAEASNGEAQFSVDLSGVDTSSVRIA 1248
pUMNK88        QYKPDGDYDANTMGVWKVRLIQQKAYNETVALTDYAEASNGEAQFSVDLSGVDTSSVRIA 1248
pCf587         QYKPDGGYDANTMGVWKVRLIQQKAYNETVALTDYAEASNGEAQFSVDLSGVDTSSVRIA 1248
               ******.*****************************************************

pKAZ3          AEAVLESPVEGYNRTELSIRPAFLTVLRGGAIGAGVEARKLSGEAPFTAVFKLSLDDRQD 1323
pMS6198A       AEAVLESPVEGYNRTELSIRPAFLTVLRGGAIGAGVEARKLSGEAPFTAVFKLSLDDRQD 1308
pKp55          AEAVLESPVEGYNRTELSIRPAFLTVLRGGAIGAGVEARKLSGEAPFTAVFKLSLDDRQD 1312
pEc19          AEAVLESPVEGYNRTELSIRPAFLTVLRGGAIGAGVEARKLSGEAPFTAVFKLSLDDRQD 1312
pAPEC1990      AEAVLESPVEGYNRTELSIRPAFLTVLRGGAIGAGVEARKLSGEAPFTAVFKLSLDDRQD 1308
pNDM-KN        AEAVLESPVEGYNRTELSIRPAFLTVLRGGAIGAGVEARKLSGEAPFTAVFKLSLDDRQD 1308
pUMNK88        AEAVLESPVEGYNRTELSIRPAFLTVLRGGAIGAGVEARKLSGEAPFTAVFKLSLDDRQD 1308
pCf587         AEAVLDSPVEGYNRTELSIRPAFLTVLRGGAIAAGVEARKLSGEAPFTAVFKLALDNRQD 1308
               *****:**************************.********************:**:***

pKAZ3          LRATGQVVWETSKDDGKTWEQFIPEDRYKYQLVKTFDKGEYQVRAKVVNVNSGAEKYTEA 1383
```

```
pMS6198A     LRATGQVVWETSKDDGKTWEQFIPEDRYKYQLVKTFDKGEYQVRAKVVNVNSGAEKYTEA 1368
pKp55        LRATGQVVWETSKDDGKTWEQFIPEDRYKYQLVKTFDKGEYQVRAKVVNVNSGAEKYTEA 1372
pEc19        LRATGQVVWETSKDDGKTWEQFIPEDRYKYQLVKTFDKGEYQVRAKVVNVNSGAEKYTEA 1372
pAPEC1990    LRATGQVVWETSKDDGKTWEQFIPEDRYKYQLVKTFDKGEYQVRAKVVNVNSGAEKYTEA 1368
pNDM-KN      LRATGQVVWETSKDDGKTWEQFIPEDRYKYQLVKTFDKGEYQVRAKVVNVNSGAEKYTEA 1368
pUMNK88      LRATGQVVWETSKDDGKTWEQFIPEDRYKYQLVKTFDKGEYQVRAKVVNVNSGAEKYTEA 1368
pCf587       LRATGQVVWETTKDDGKTWEQFIPEERYKYQLVKTFDKGEYQVRAKVVNVNSGAEKYTEA 1368
             ***********:*************:**********************************

pKAZ3        VSVVAYDKPDIAVIGPTTLFVGSEGKYTANLTLNDEPISGGNAIVEWSTDGGKTYAQTGD 1443
pMS6198A     VSVVAYDKPDIAVIGPTTLFVGSEGKYTANLTLNDEPISGGNAIVEWSTDGGKTYAQTGD 1428
pKp55        VSVVAYDKPDIAVIGPTTLFVGSEGKYTANLTLNDEPISGGNAIVEWSTDGGKTYAQTGD 1432
pEc19        VSVVAYDKPDIAVIGPTTLFVGSEGKYTANLTLNDEPISGGNAIVEWSTDGGKTYAQTGD 1432
pAPEC1990    VSVVAYDKPDIAVIGPTTLFVGSEGKYTANLTLNDEPISGGNAIVEWSTDGGKTYAQTGD 1428
pNDM-KN      VSVVAYDKPDIAVIGPTTLFVGSEGKYTANLTLNDEPISGGNAIVEWSTDGGKTYAQTGD 1428
pUMNK88      VSVVAYDKPDIAVIGPTTLFVGSEGKYTANLTLNDEPISGGNAIVEWSTDGGKTYAQTGD 1428
pCf587       VSVVAYDKPDIAVVGPTTLFVGSEGKYTASLALNDEPITDGNAVVEWSTDGGKTYSHKGN 1428
             *************:***************.*:******:.***:***********::.*:

pKAZ3        SITLSSDEETRYRLWARVRSATAPADDGYAYEVAKTAVDFRAVKAPRPYVTGPRVIETGK 1503
pMS6198A     SITLSSDEETRYRLWARVRSATAPADDGYAYEVAKTAVDFRAVKAPRPYVTGPRVIETGK 1488
pKp55        SITLSSDEETRYRLWARVRSATAPADDGYAYEVAKTAVDFRAVKAPRPYVTGPRVIETGK 1492
pEc19        SITLSSDEETRYRLWARVRSATAPADDGYAYEVAKTAVDFRAVKAPRPYVTGPRVIETGK 1492
pAPEC1990    SITLSSDEETRYRLWARVRSATAPADDGYAYEVAKTAVDFRAVKAPRPYVTGPRVIETGK 1488
pNDM-KN      SITLSSDEETRYRLWARVRSATAPADDGYAYEVAKTAVDFRAVKAPRPYVTGPRVIETGK 1488
pUMNK88      SITLSSDEETRYRLWARVRSATAPADDGYAYEVAKTAVDFRAVKAPRPYVTGPRVIETGK 1488
pCf587       SITLSSDEETRYRLWARVRSANAPADDDYAYDVAKTAVDFRAVKAPRPYVVGPQVIETGK 1488
             *********************.*****.***:******************.**:******

pKAZ3        KYVFKAETSLPYRGMDVKLNGFFTLPDGSIVQGDTAEYVPSDTDLNQATVETKYTTWIEG 1563
pMS6198A     KYVFKAETSLPYRGMDVKLNGFFTLPDGSIVQGDTAEYEPSDTDLNQATVETKYTTWIEG 1548
pKp55        KYVFKAETSLPYRGMDVKLNGFFTLPDGSIVQGDTAEYEPSDTDLNQATVETKYTTWIEG 1552
pEc19        KYVFKAETSLPYRGMDVKLNGFFTLPDGSIVQGDTAEYEPSDTDLNQATVETKYTTWIEG 1552
pAPEC1990    KYVFKAETSLPYRGMDVKLNGFFTLPDGSIVQGDTAEYEPSDTDLNQATVETKYTTWIEG 1548
pNDM-KN      KYVFKAETSLPYRGMDVKLNGFFTLPDGSIVQGDTAEYEPSDTDLNQATVETKYTTWIEG 1548
pUMNK88      KYVFKAETSLPYRGMDVKLNGFFTLPDGSIVQGDTAEYEPSDTDLNQATVETKYTTWIEG 1548
pCf587       KYVFSAKTSLPYRGMDVKLNGFFTLPDGSIVEGDTAEYVPSDNDLNQATVETKYTTWIEG 1548
             ****.*:************************:****** ***.*****************

pKAZ3        YRDQGAEASHSLRSRVWQYVWPSFGMQVRKNADVAPATITASVRPIAFNGKLEEPTYEWE 1623
pMS6198A     YRDQGAEASHSLRSRVWQYVWPSFGMQVRKNADVAPATITASVRPIAFNGKLEEPTYEWE 1608
pKp55        YRDQGAEASHSLRSRVWQYVWPSFGMQVRKNADVAPATITASVRPIAFNGKLEEPTYEWE 1612
pEc19        YRDQGAEASHSLRSRVWQYVWPSFGMQVRKNADVAPATITASVRPIAFNGKLEEPTYEWE 1612
pAPEC1990    YRDQGAEASHSLRSRVWQYVWPSFGMQVRKNADVAPATITASVRPIAFNGKLEEPTYEWE 1608
pNDM-KN      YRDQGAEASHSLRSRVWQYVWPSFGMQVRKNADVAPATITASVRPIAFNGKLEEPTYEWE 1608
pUMNK88      YRDQGAEASHSLRSRVWQYVWPSFGMQVRKNADVAPATITASVRPIAFNGKLEEPTYEWE 1608
pCf587       YRDKGAEASHSLRSRVWQYVWPGFAMQVRKSADAAPATITASVRPIAFNGKLEEPTYEWE 1608
             ***:******************.*.*****.**.**************************

pKAZ3        LPEGAVIQDQRQDIVRSFVINEPGDYNIKVTVRDARGHETVIEQPLKIGQAEPYAIDLQY 1683
pMS6198A     LPEGAVIQDQRQDIVRSFVINEPGDYNIKVTVRDARGHETVIEQPLKIGQAAPYAIDLQY 1668
pKp55        LPEGAVIQDQRQDIVRSFVINEPGDYNIKVTVRDARGHETVIEQPLKIGQAAPYAIDLQY 1672
pEc19        LPEGAVIQDQRQDIVRSFVINEPGDYNIKVTVRDARGHETVIEQPLKIGQAAPYAIDLQY 1672
pAPEC1990    LPEGAVIQDQRQDIVRSFVINEPGDYNIKVTVRDARGHETVIEQPLKIGQAAPYAIDLQY 1668
pNDM-KN      LPEGAVIQDQRQDIVRSFVINEPGDYNIKVTVRDARGHETVIEQPLKIGQAAPYAIDLQY 1668
pUMNK88      LPEGAVIQDQRQDIVRSFVINEPGDYNIKVTVRDARGHETVIEQPLKIGQAAPYAIDLQY 1668
pCf587       LPEGAVIQDKKQDIVRSFVVNEPGDYNIKVTVRDARGHEAVIEKSFKIDQSKPYAIDLQY 1668
             *********::********:*******************:***: :**.*: ********

pKAZ3        SGSNKYEREPLDVLLRPYISGGHPRDRISTRVYSVDGTPLESSGYYGRATLGAGEHSIKL 1743
pMS6198A     SGSNKYEREPLDVLLRPYISGGHPRDRISTRVYSVDGTPLESSGYYGRATLGAGEHSIKL 1728
pKp55        SGSNKYEREPLDVLLRPYISGGHPRDRISTRVYSVDGTPLESSGYYGRATLGAGEHSIKL 1732
pEc19        SGSNKYEREPLDVLLRPYISGGHPRDRISTRVYSVDGTPLESSGYYGRATLGAGEHSIKL 1732
pAPEC1990    SGSNKYEREPLDVLLRPYISGGHPRDRISTRVYSVDGTPLESSGYYGRATLGAGEHSIKL 1728
pNDM-KN      SGSNKYEREPLDVLLRPYISGGHPRDRISTRVYSVDGTPLESSGYYGRATLGAGEHSIKL 1728
pUMNK88      SGSNKYEREPLDVLLRPYISGGHPRDRISTRVYSVDGTPLESSGYYGRATLGAGEHSIKL 1728
pCf587       SGSNKYEREPLDVLLRPYISGGHPRDRILTHVYSVDGTPLENSGYYGKATLRAGEHSIKL 1728
             **************************** *:**********.*****:*** ********

pKAZ3        KITSEMGHEAEGEVNINVAENKLPACSLSSRETVGSWIVYANCEDTDGRMKSYEWTIAGE 1803
pMS6198A     KITSEMGHEAEGEVNINVAENKLPACSLSSRETVGSWIVYANCEDTDGRMKSYEWTIAGE 1788
pKp55        KITSEMGHEAEGEVNINVAENKLPACSLSSRETVGSWIVYANCEDTDGRMKSYEWTIAGE 1792
pEc19        KITSEMGHEAEGEVNINVAENKLPACSLSSRETVGSWIVYANCEDTDGRMKSYEWTIAGE 1792
pAPEC1990    KITSEMGHEAEGEVNINVAENKLPACSLSSRETVGSWIVYANCEDTDGRMKSYEWTIAGE 1788
pNDM-KN      KITSEMGHEAEGEVNINVAENKLPACSLSSRETVGSWIVYANCEDTDGRMKSYEWTIAGE 1788
pUMNK88      KITSEMGHEAEGEVNINVAENKLPACSLSSRETVGSWIVYANCEDTDGRMKSYEWTIAGE 1788
pCf587       KITSEMGHEAEGEVSINVAENKLPVCSLKSRETVGSWIVYANCEDTDGRMKSYEWTIAGE 1788
```

```
                **************.*********.***.********************************

pKAZ3           LQSISSDRVTISKGTYETMPTISLVGVDDSGGKSEAVTMN  1843
pMS6198A        LQSISSDRVTISKGTYETMPTISLVGVDDSGGKSEAVTMN  1828
pKp55           LQSISSDRVTISKGTYETMPTISLVGVDDSGGKSEAVTMN  1832
pEc19           LQSISSDRVTISKGTYETMPTISLVGVDDSGGKSEAVTMN  1832
pAPEC1990       LQSISSDRVTISKGTYETMPTISLVGVDDSGGKSEAVTMN  1828
pNDM-KN         LQSISSDRVTISKGTYETMPTISLVGVDDSGGKSEAVTMN  1828
pUMNK88         LQSISSDRVTISKGTYETMPTISLVGVDDSGGKSEAVTMN  1828
pCf587          LKSISSDRVTISKGPNKTMPTISLVGVDDSGGRSEAVTMN  1828
                *:************  :***************:*******
```

**Figure S2.** Alignment of proteins containing Big domains in IncA/C plasmids. Big 3_2 and Big3_3 domains are highlighted in blue and green boxes, respectively.

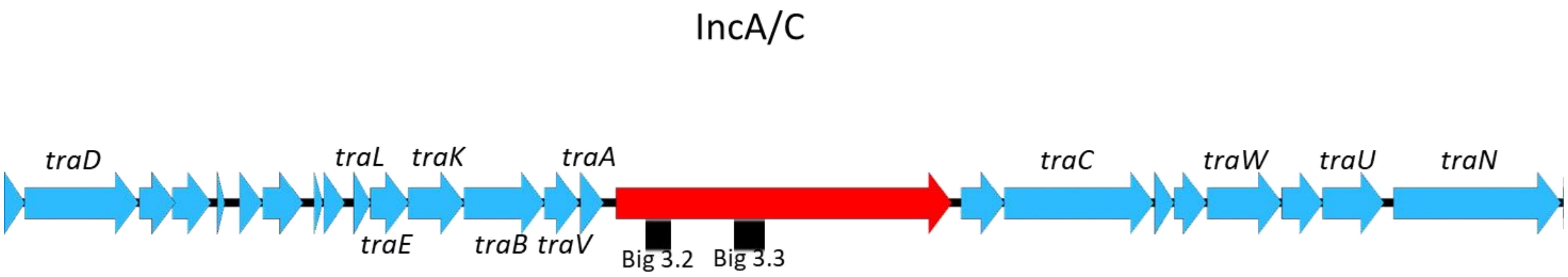


**Figure S3.** Genomic context of the identified Big proteins (in red) in IncA/C plasmids. The black boxes correspond to the Big 3.2 and Big 3.3 domains.

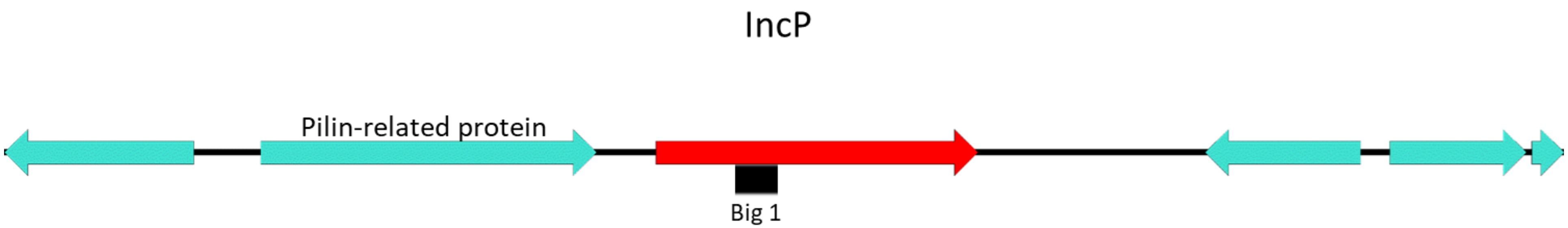


**Figure S4.** Genomic context of the identified Big proteins (in red) in IncP2 plasmids. The black box corresponds to the Big 1 domain.

```
CKA44          --------------------------------------------------------MGLT     4
HMPREF1223     MFANLKALAVAGAFFLMSVTTTVSADSWKPIRSGSSSSGWQKVVCDRSGNGWRSCNMGLT     60
pOZ176         --------------------------------------------------------MGLT     4
pJB37          MFANLKALAVAGAFFLMSVTTTVSADSWKPIRSGSSSSGWQKVVCDRSGNGWRSCNMGLT     60
pPWIS1         --------------------------------------------------------MGLT     4
pAPA25         --------------------------------------------------------MGLT     4
pNK546KPC      MFANLKALAVAGAFFLMSVTTTVSADSWKPIRSGSSSSGWQKVVCDRSGNGWRSCNMGLT     60
RN02           --------------------------------------------------------MGLT     4
                                                                       ****

CKA44          IVIQATPGSLAALGEKATLVATVQDYDGNNAGRGVVINWTTSDGGLSAATTTTDANGQTS     64
HMPREF1223     IVIQATPGSLAALGEKATLVATVQDYDGNNAGRGVVINWTTSDGGLSAATTTTDANGQTS     120
pOZ176         IVIQATPGSLAALGEKATLVATVQDYDGNNAGRGVVINWTTSDGGLSAATTTTDANGQTS     64
pJB37          IVIQATPGSLAALGEKATLVATVQDYDGNNAGRGVVINWTTSDGGLSAATTTTDANGQTS     120
pPWIS1         IVIQATPGSLAALGEKATLVATVQDYDGNNAGRGVVINWTTSDGGLSAATTTTDANGQTS     64
pAPA25         IVIQATPGSLAALGEKATLVATVQDYDGNNAGRGVVINWTTSDGGLSAATTTTDANGQTS     64
pNK546KPC      IVIQATPGSLAALGEKATLVATVQDYDGNNAGRGVVINWTTSDGGLSAATTTTDANGQTS     120
RN02           IVIQATPGSLAALGEKATLVATVQDYDGNNAGRGVVINWTTSDGGLSAATTTTDANGQTS     64
               ************************************************************

CKA44          VVLTSSKTIGGATVSATSPAEGGTGQITVPFTDKWVSTSAMYSAWQDSGAPYSCSAWSPD     124
HMPREF1223     VVLTSSKTIGGATVSATSPAEGGTGQITVPFTDKWVSTSAIYSAWQDSGAPYSCSAWSPD     180
pOZ176         VVLTSSKTIGGATVSATSPAEGGTGQITVPFTDKWVSTSAMYSAWQDSGAPYSCSAWSPD     124
pJB37          VVLTSSKTIGGATVSATSPAEGGTGQITVPFTDKWVSTSAMYSAWQDSGAPYSCSAWSPD     180
pPWIS1         VVLTSSKTIGGATVSATSPAEGGTGQITVPFTDKWVSTSAMYSAWQDSGAPYSCSAWSPD     124
pAPA25         VVLTSSKTIGGATVSATSPAEGGTGQITVPFTDKWVSTSAMYSAWQDSGAPYSCSAWSPD     124
pNK546KPC      VVLTSSKTIGGATVSATSPAEGGTGQITVPFTDKWVSTSAMYSAWQDSGAPYSCSAWSPD     180
RN02           VVLTSSKTIGGATVSATSPAEGGTGQITVPFTDKWVSTSAMYSAWQDSGAPYSCSAWSPD     124
               ****************************************:*******************

CKA44          VSTINQGTSFTQSAVCYQNQIAYQQNREVSLVTGQVRNVGGVIPLYQTVQAARSQQAVGT     184
HMPREF1223     ASTINQGTSFTQSAVCYQNQIAYQQNREVSLVTGQVRNVGGVIPLYQTVQAARSQQAVGT     240
pOZ176         ASTINQGTSFTQSAVCYQNQIAYQQNREVSLVTGQVRNVGGVIPLYQTVQAARSQQAVGT     184
pJB37          ASTINQGTSFTQSAVCYQNQIAYQQNREVSLVTGQVRNVGGVIPLYQTVQAARSQQAVGT     240
pPWIS1         ASTINQGTSFTQSAVCYQNQIAYQQNREVSLVTGQVRNVGGVIPLYQTVQAARSQQAVGT     184
pAPA25         ASTINQGTSFTQSAVCYQNQIAYQQNREVSLVTGQVRNVGGVIPLYQTVQAARSQQAVGT     184
pNK546KPC      ASTINQGTSFTQSAVCYQNQIAYQQNREVSLVTGQVRNVGGVIPLYQTVQAARSQQAVGT     240
RN02           ASTINQGTSFTQSAVCYQNQIAYQQNREVSLVTGQVRNVGGVIPLYQTVQAARSQQAVGT     184
               .***********************************************************

CKA44          KQSTPSCAWSSFTKNGVYATGWDHGVSNTGGPKQGYRLYLGQYIGEVANATDSFAYNGRI     244
HMPREF1223     KQSTPSCAWSSFTKNGVYATGWDHGVSNTGGPKQGYRLYLGQYIGEVANATDSFAYNGRI     300
pOZ176         KQSTPSCAWSSFTKNGVYATGWDHGVSNTGGPKQGYRLYLGQYIGEVANATDSFAYNGRI     244
pJB37          KQSTPSCAWSSFTKNGVYATGWDHGVSNTGGPKQGYRLYLGQYIGEVANATDSFAYNGRI     300
pPWIS1         KQSTPSCAWSSFTKNGVYATGWDHGVSNTGGPKQGYRLYLGQYIGEVANATDSFAYNGRI     244
pAPA25         KQSTPSCAWSSFTKNGVYATGWDHGVSNTGGPKQGYRLYLGQYIGEVANATDSFAYNGRI     244
pNK546KPC      KQSTPSCAWSSFTKNGVYATGWDHGVSNTGGPKQGYRLYLGQYIGEVANATDSFAYNGRI     300
RN02           KQSTPSCAWSSFTKNGVYATGWDHGVSNTGGPKQGYRLYLGQYIGEVANATDSFAYNGRI     244
               ************************************************************

CKA44          YTIGKFRQSTCLGKNCASSREEYEACSVPQ     274
HMPREF1223     YTIGKFRQSTCLGKNCASSREEYEACSVPQ     330
pOZ176         YTIGKFRQSTCLGKNCASSREEYEACSVPQ     274
pJB37          YTIGKFRQSTCLGKNCASSREEYEACSVPQ     330
pPWIS1         YTIGKFRQSTCLGKNCASSREEYEACSVPQ     274
pAPA25         YTIGKFRQSTCLGKNCASSREEYEACSVPQ     274
pNK546KPC      YTIGKFRQSTCLGKNCASSREEYEACSVPQ     330
RN02           YTIGKFRQSTCLGKNCASSREEYEACSVPQ     274
               ******************************
```

**Figure S5.** Alignment of proteins containing Big domain in IncP2 plasmids. Big_1 domain is highlighted in orange.

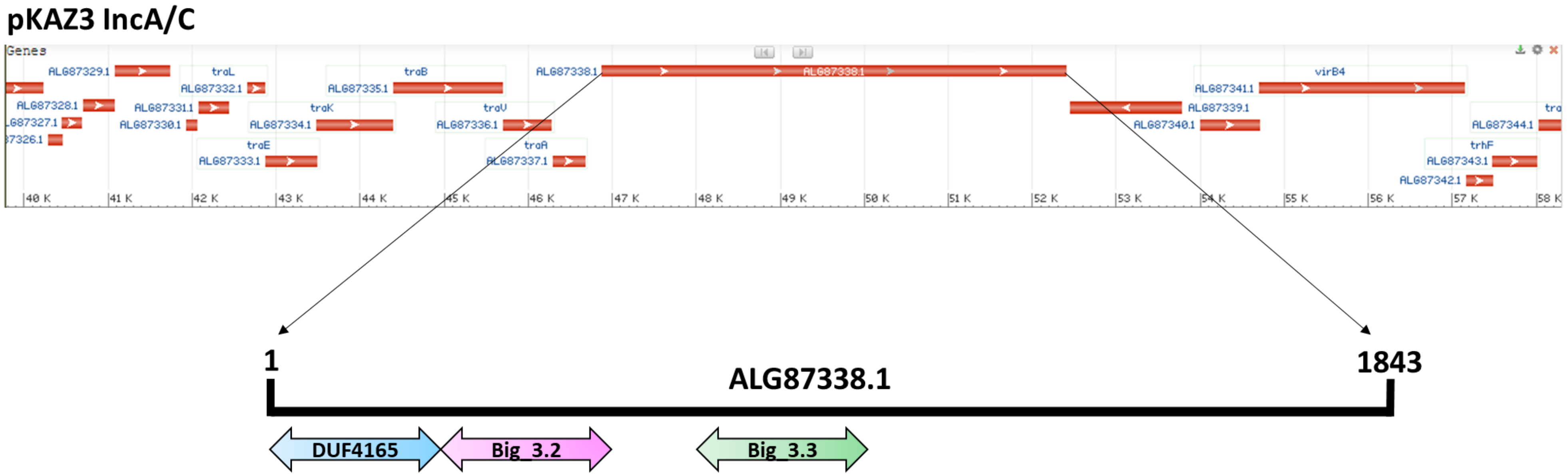


**Figure S6.** Genomic context of the identified Big protein (ALG87338.1) in the IncA/C plasmid pKAZ3. The ALG87338.1 protein is an 1843 AA protein and contains a DUF4165, Big_3.2 and Big_3.3 domains.

| Strains | Relevant characteristics | Reference/Source |
|---|---|---|
| SL1344 | *rspL, hisG* | [1] |
| SL1344ibplac | *ibpA*$_{420}$*::lacZ.* Km$^r$ | [2] |
| SL1344 Δ*fliC* Δ*fljB* | *fliC*::FRT *fljB*::FRT | This work |
| **Plasmids** | **Relevant characteristics** | **Reference/Source** |
| R27 | IncHI1, Tc$^r$ | [3] |
| pKAZ3 | IncA/C, Cb$^r$, Tcr | Dr. Álvaro San Millán |
| R27 Δ*rsp2* | R27 *rsp2*::FRT, Tc$^r$ | This work |
| R27 RSP-Flag | R27 *rsp*::Flag, Tc$^r$, Km$^r$ | [4] |
| R27 RSP2-Flag | R27 *rsp2*::Flag, Tc$^r$, Km$^r$ | This work |
| R27 Δ*rsp* | R27 *rsp*::FRT, Tc$^r$ | [4] |
| R27 Δ*rsp* RSP2-Flag | R27 *rsp*::FRT, *rsp2*::Flag, Tc$^r$, Km$^r$ | This work |
| R27 Δ*trhC* | R27 *trhC*::FRT, Tc$^r$ | [4] |
| R27 Δ*trhC* RSP-Flag | R27 *trhC*::Cm, *rsp*::Flag. Cm$^r$, Km$^r$, Tc$^r$ | [4] |
| R27 Δ*trhC* RSP2-Flag | R27 *trhC*::Cm, *rsp2*::Flag. Cm$^r$, Km$^r$, Tc$^r$ | This work |
| R27 Δ*trhH* | R27 *trhH*::FRT, Tc$^r$ | This work |
| R27 Δ*trhH* RSP-Flag | R27 *trhH*::FRT, *rsp*::Flag. Km$^r$, Tc$^r$ | This work |
| R27 Δ*trhH* RSP2-Flag | R27 *trhH*::FRT, *rsp2*::Flag. Km$^r$, Tc$^r$ | This work |
| R27 Δ*trhA* | R27 *trhA*::FRT, Tc$^r$ | This work |

| | | |
|---|---|---|
| R27 Δ*trhA* RSP-Flag | R27 *trhA*::FRT, *rsp*::Flag. Km$^r$, Tc$^r$ | This work |
| R27 Δ*trhA* RSP2-Flag | R27 *trhA*::FRT, *rsp2*::Flag. Km$^r$, Tc$^r$ | This work |
| pLG338-*rsp* | pLG338-30 + *rsp* from R27 | [4] |
| pLG338-30 | ori$_p$SC101, Cb$^r$ | [5] |
| pLG338-*rsp2* | pLG338-30 + *rsp2* from R27 | This work |
| pBR322-*trhC* | pBR322 + *trhC* from R27 | [4] |
| pKD4 | *bla* FRT *ahp* FRT PS1 PS2 oriR6K Km$^r$, Cb$^r$ | [6] |
| pKD3 | *bla* FRT *cat* FRT PS1 PS2 oriR6K Cm$^r$, Cb$^r$ | [6] |
| pSUB11 | Flag- and Km$^r$-coding template vector | [7] |
| pKD46 | *oriR101, repA101 (ts), AraBp-gam-bet-exo* | [6] |
| pKD46-Km$^R$ | *oriR101, repA101 (ts), AraBp-gam-bet-exo*, Km$^r$ | This work |
| pKAZ3 ΔALG87338.1 | pKAZ3 ALG87338.1:: Cm, Cm$^r$, Tc$^r$ | This work |

**Table S3.** Bacterial strains and plasmids used in this work.

| Name | Sequence 5’ - 3’ | Use |
|---|---|---|
| **fliC_SL1344_P1** | CATCAAGTTGTAATTGATAAGGAAAAGATCATGGCA GTGTAGGCTGGAGCTGCTTC | ***fliC* deletion** |
| **fliC_SL1344_P2** | GTACCACGTGTCGGTGAATCAATCGCCGGATTAACG CATATGAATATCCTCCTTAGT | ***fliC* deletion** |
| **fliC_SL1344_P1Up** | CGTTCTTTGTCAGGTCTGTCAACAACTGG | ***fliC* deletion confirmation** |
| **fliC_SL1344_P2down** | GCAGGCAAGACTCAGAGAGTTACGCACC | ***fliC* deletion confirmation** |
| **fljB_SL1344_P1** | GCTTTATCAAAAACCTTCCAAAAGGAAAATTTTATG GCAGTGTAGGCTGGAGCTGCTTC | ***fljB* deletion** |
| **fljB_SL1344_P2** | AATTCACGGGGCTGAATAAAACGAAATAAATTAAC GCATATGAATATCCTCCTTAGT | ***fljB* deletion** |
| **fljB_SL1344_P1up** | CGCCACCAGGTTTTTCACGCT | ***fljB* deletion confirmation** |
| **fljB_SL1344_P2down** | CCTGTCGTTTTGCCAGTCAAAACCTGTCC | ***fljB* deletion confirmation** |
| **RSP2_R27_P1** | GGTAAAATTTTCTGGCCAACCAGGAGAACCGAAAT GAAATTTGTGTAGGCTGGAGCTGCTTC | ***rsp2* deletion** |
| **RSP2_R27_P2** | AAGCCCCGTAATACGGGGCCGGTTCGGAGGCAGTT ACTGGATCATATGAATATCCTCCTTAGT | ***rsp2* deletion** |
| **RSP2_R27_P1up** | CAGGCCTGCCGATTAAATCTG | ***rsp2* deletion confirmation** |
| **RSP2_R27_P2down** | CTGGCTCTGCATTCGAATTTC | ***rsp2* deletion confirmation** |
| **RSP2_R27_3xP1** | CGGTGAATCAAGCTTCATGATCGAGCTGCCGCAAAT CCAGGACTACAAAGACCATGACGG | **RSP2-Flag** |

| | | |
|---|---|---|
| **RSP2_R27_3xP2** | AAGCCCCGTAATACGGGGCCGGTTCGGAGGCAGTT ACTGGCATATGAATATCCTCCTTAG | **RSP2-Flag** |
| **RSP2_R27_3xP1up** | GCGGTAGATGAAGCCGGTAAT | **RSP2-Flag confirmation** |
| **RSP2_R27_3xP2down** | CTGGCTCTGCATTCGAATTTC | **RSP2-Flag confirmation** |
| **R27_RSP2_pLG EcoRI fw** | CGGAATTCCTACATCTGCGTCGTACTGAA | ***rsp2* cloning** |
| **R27_RSP2_pLG BamHI rv** | CGGGATCCTGCGACTAAATCAGCCTGTTT | ***rsp2* cloning** |
| **trhH_R27_P1** | ACTATCTGAATGTGGGTGGAAGTTTAGGAGGTGCA TATGCGCGTGTAGGCTGGAGCTGCTTC | ***trhH* deletion** |
| **trhH_R27_P2** | CGCCAACTGTATAGATGTTGTAATCCATATCAGACC TCAGTTCATATGAATATCCTCCTTAGT | ***trhH* deletion** |
| **trhH_R27_P1up** | ACCATTGGCGAGGATGGCGTA | ***trhH* deletion confirmation** |
| **trhH_R27_P2down** | GGCTGGCTCCAGCCAGTACCG | ***trhH* deletion confirmation** |
| **trhA_R27_P1** | TGCGTTTTCGTGAATTCAAATCAACACGGAGTAATT ATGGAAGTGTAGGCTGGAGCTGCTTC | ***trhA* deletion** |
| **trhA_R27_P2** | GGGGAATATCCCCCTCATTGTTATTGTTCTAACAAAT CACAGCATATGAATATCCTCCTTAGT | ***trhA* deletion** |
| **trhA_R27_P1up** | GGATGAATGCCATAAAAATGG | ***trhA* deletion confirmation** |
| **trhA_R27_P2down** | CCGCTCGCGATAGTCACGAAT | ***trhA* deletion confirmation** |
| **KT** | CGGCCACAGTCGATGAATCC | **Confirmation $Km^R$ insertion** |
| **CatC1** | TTATACGCAAGGCGACAAGG | **Confirmation $Cm^R$ insertion** |

| | | |
|---|---|---|
| **PkmXmnIFw** | GAAACGTTTCAGCACTCAGGGCGCAAGGGCT | **Amplification $Km^r$ for cloning into pKD46** |
| **PkmXmnIRv** | GAAACGTTTCTCAGAAGAACTCGTCAAGAAG | **Amplification $Km^r$ for cloning into pKD46** |
| **ALG87338.1P1** | TTGGGTTCTTGTAACTCAAAACGAATGGAGAATGCG ATGATCGTGTAGGCTGGAGCTGCTTC | ***ALG87338.1* deletion** |
| **ALG87338.1P2** | ATTGCGGTCTGAACCGGGCTTTGAGAGGAATGAAC CTTAGTTCATATGAATATCCTCCTTAGT3 | ***ALG87338.1* deletion** |
| **ALG87338.1P1up** | ACGGGTGATAGGGGCAGCCTA | ***ALG87338.1* deletion confirmation** |
| **ALG87338.1P2down** | ATGCCATCACACTTCGCAGC | ***ALG87338.1* deletion confirmation** |

**Table S4.** Oligonucleotides used in this study.
